# Molecular dynamics driving phenotypic divergence among KRAS mutants in pancreatic tumorigenesis

**DOI:** 10.1101/2025.05.28.656689

**Authors:** Adrien Grimont, David J. Falvo, Whitney J. Sisso, Paul Zumbo, Christopher W. Chan, Francisco Santos, Grace Pan, Megan Cleveland, Tomer M. Yaron, Alexa Osterhoudt, Yinuo Meng, Maria Paz Zafra, William B. Fall, Andre F. Rendeiro, Erika Hissong, Rhonda K. Yantiss, Doron Betel, Mark A. Magnuson, Steven D. Leach, Anil K. Rustgi, Lukas E. Dow, Rohit Chandwani

## Abstract

Inflammation in the pancreas drives acinar-to-ductal metaplasia (ADM), a progenitor-like state that can be hijacked by mutant ***Kras*** in the formation of pancreatic cancer (PDAC). How these cell fate decisions vary according to KRAS mutation remains poorly understood. To define mutation-specific lineage reversion and tumor initiation, we implement novel Ptf1a-TdTomato mice and multiple KRAS mutants across an array of genetic, pharmacologic, and inflammatory perturbations ***in vivo***. Whereas KRAS^G12D^ co-opts injury to enable lineage reversion, enhancer reprogramming, and tumor initiation, KRAS^G12R/V^ can initiate but not sustain dedifferentiated and neoplastic transcriptional and epigenetic programs. We find the KRAS^G12R/V^ defects consist of a failure to invoke robust EGFR signaling and activate Rac1/Vav1, with constitutive Akt activation ***in vivo*** sufficient to rescue the tumorigenic potential of KRAS^G12R^. As the marked heterogeneity among KRAS variants begins early in tumorigenesis, these data are crucial to understanding mutation-specific oncogenic trajectories and directing the implementation of ***KRAS***-directed therapeutics.

**SIGNIFICANCE:** Defining how *KRAS* mutants drive distinct outcomes in human pancreatic cancer is critical for developing allele-specific therapeutic approaches. This study unveils a hierarchy among KRAS^G12D^, KRAS^G12V^, and KRAS^G12R^ to drive tumor initiation, owing to heterogeneous activation of EGFR, PI3K/AKT, and RAC1 signaling, thus revealing mutation-specific evolutionary paths in pancreatic tumorigenesis.

## INTRODUCTION

Living systems exist in an apparent contradiction – they ensure their stability through a profound capacity for change. Indeed, in the pancreas, the cellular plasticity of the acinar compartment – namely to undergo acinar-ductal metaplasia (ADM) and regenerate in the context of inflammatory injury – is essential to tissue homeostasis.^1–3^ Multiple reports have suggested that ADM is not a *bona fide* transdifferentiation event, but one that features re-expression of pancreatic progenitor markers, including Nestin, KLF4, and KLF5^1,4,5^. Indeed, reprogramming of the acinar compartment via expression of the pluripotency factors OCT4, SOX2, KLF4, and cMYC (OSKM) can drive tumorigenesis in concert with mutant *Kras* in lieu of inflammatory injury induced by the cholecystokinin analog caerulein.^6^ The extent to which lineage reversion – apart from expression of these selected markers – is a conserved feature of inflammatory injury or of neoplastic cell fates has not been fully articulated.

Nevertheless, the metaplastic state is key, largely because it is this cell fate that is in turn hijacked by mutant *Kras* in the development of pancreatic intraepithelial neoplasia (PanIN) and pancreatic ductal adenocarcinoma (PDAC).^7–11^ Preclinical models have demonstrated sufficiency of mutant *Kras* and p53 to recapitulate the full spectrum of human disease from tumor initiation to metastasis.^12^ In the adult mouse, however, induction of ADM via inflammation is essential, and it is the synergy between mutant *Kras*^G12D^ and inflammation to rewire chromatin that drives neoplastic commitment.^7,8,10^Indeed, our group and others have shown that metaplasia does not fully resolve even over prolonged periods, and continues to support tumorigenesis despite temporal separation of inflammatory and oncogenic *Kras*^G12D^ activation^13–14^, highlighting the instrumental role of injury in driving the lineage plasticity essential for neoplastic commitment.

Mutant *KRAS*, found in 95% of PDAC, is essential to both tumor initiation and maintenance^15–16^, with *KRAS*^G12D^ the most common (35% of patients) and well-studied of the mutant alleles. Similar to *KRAS*^G12D^, *KRAS*^G12V^, occurring in 30% of patients, also requires in acinar-cell specific models the presence of an inflammatory insult to drive tumor formation^7^, but the degree to which epigenetic reprogramming is recapitulated in this driver mutational context has not been explored. Finally, early studies suggest that KRAS^G12R^, which occurs in 15% of PDAC, is diminished relative to KRAS^G12D^ in its oncogenic potential.^17–18^ A thorough comparative analysis of the three most common mutations in KRAS – G12D, G12V, and G12R – has not been performed, such that it remains unknown whether these are functionally equivalent in their capacity to drive oncogenic chromatin remodeling or to yield similar or disparate phenotypic outcomes, particularly in physiologic models of adult mutant *Kras* activation. As the precise *KRAS* mutation has been recently shown to powerfully influence clinical outcome and tumor features in human disease^19–20^, the distinct biology of the *Kras* mutants is imperative to articulate, especially in the context of emergent approaches to target mutant KRAS.^21–22^

Herein we evaluated the molecular dynamics in injury and across the three most common mutant *Kras* alleles (G12D, G12V, and G12R) in tumor initiation, leveraging in each lineage-tracing models of adult *Kras* activation to carefully interrogate specific cellular compartments. Across all the *Kras* mutants, we find little evidence of an ability of the oncogene to alter chromatin on its own. However, when inflammation is superimposed, *Kras*^G12D^ can robustly rewire chromatin to drive accumulation of AP-1 chromatin accessibility along with lineage reversion to resemble early progenitors; *Kras*^G12R^, stalled in ADM, does not initiate the key cell-intrinsic epigenetic progenitor programs, with *Kras*^G12V^ also reduced in oncogenic potential as compared to *Kras*^G12D^. Surprisingly, we observe divergence among the mutants not in KRAS activity nor in oncogenic MAPK signaling but in EGFR signaling and RAC1 activation. Finally, systematic perturbation of the oncogenic signals downstream of KRAS showed a sufficiency of constitutive activation of PI3K/AKT signaling, but not of RAC1 activation nor enhanced MAPK signaling nor even biallelic disruption of p53, to drive tumor initiation in KRAS^G12R^. Together, our data reveal the molecular basis of divergence among the *Kras* mutants, localize the phenotypic separation as occurring not with lineage plasticity but in neoplastic commitment, and establish the presence of mutation-specific oncogenic trajectories in pancreatic cancer.

## RESULTS

### Injury induces lineage reversion to a dedifferentiated cell state

Acinar metaplasia in response to damage is traditionally explained as a transition toward a ductal lineage, as lineage-traced metaplastic lesions adopt a ductal morphology while also expressing a handful of ductal-specific markers.^1–3^ Metaplastic lesions have been demonstrated to contain cells that also express markers associated with pancreatic progenitor cells (e.g., Pdx1, Klf4, Klf5, Nestin).^1,3–5,23^ To define the extent to which acinar cells transdifferentiate to ductal cells or whether they adopt an alternative cell state (i.e., dedifferentiate toward pancreatic progenitor cells), we sought to profile the transcriptional and epigenetic landscapes of normal acinar cells, ductal cells, early and late pancreatic progenitor cells, as well as acinar-derived metaplastic cells. To this end, we first generated Ptf1a-tdTomato (Ptf1a-tdT) mice to carry a knock-in allele containing both the tdTomato reporter and a functional copy of the Pt1fa transcription factor, which safeguards Ptf1a’s endogenous expression levels to ensure normal pancreatic development (Fig. S1A). Whole-organ fluorescent imaging demonstrated normal gross pancreatic morphology and tdTomato expression in both heterozygous (Ptf1a^tdT/+^) and homozygous (Ptf1a^tdT/tdT^) adult mice (Fig. S1B). In addition, we confirmed exclusive labeling of the acinar compartment by immunofluorescent (IF) staining of tdTomato and GFP in a collagenase digested pancreas isolated from an adult Ptf1a^tdT/+^; Pdx1^GFP/+^ mouse, as well as by IF on adult mouse pancreas sections (Fig. S1C-D).

Using these mice, we sought to molecularly characterize progenitor compartments during pancreatic development. Embryos from pregnant Ptf1a-tdTomato (Ptf1a-tdT) mice were dissected at embryonic day 10.5 (e10.5, early progenitors) and embryonic day 15.5 (e15.5, late progenitors), dissociated, and subjected to flow cytometry (Fig. S1E). Employing a different approach, we sorted ductal cells from acinar cells derived from C57BL/6 mice on the basis of the cell surface markers Cd49F and Cd133 (Fig. S1E). Validation of adult acinar and ductal cell sorting was performed via RT-qPCR on representative marker genes, confirming successful delineation of the exocrine cell types (Fig. S1F). We performed RNA-seq on each of the above sorted populations (early progenitor, late progenitor, ductal cells, and acinar cells [the latter sorted from *Mist1*-Cre^ERT2^; CAGs-LSL-tdTomato (MT) mice]) to characterize their transcriptional profiles. Transcriptionally, early progenitors were most distinct, whilst late progenitors, an intermediate cell fate, were most like the other cell types (Fig. S1G). These data were integrated to generate gene sets corresponding to each of the cell types, as well as to ‘exocrine’ and ‘progenitor’ cellular states (Fig. S2A-E).

To evaluate the molecular changes elicited with pancreatitis, we turned to MT mice treated with caerulein to induce pancreatitis (Fig. S1H) As expected, caerulein-treated mice showed signs of severe tissue damage with both significant stromal activation and advanced metaplastic lesions, while saline-treated mice show normal tissue architecture and no signs of damage (Fig. S1H). Sorted tdT+ cells were subjected to RNA-sequencing. Principal component analysis (PCA) demonstrated ADM largely undergoes a ductal transition (Fig. S1I), with short Euclidean distances to both ductal and late progenitor types (Fig. S1J). Using the baseline molecular profiles generated from the normal cell types, we assessed how ADM transcriptionally relates to differentiated and progenitor cell ‘benchmarks’. Indeed, using Gene Set Enrichment Analysis (GSEA), we observed that ADM featured a robust enrichment of our progenitor and specifically late progenitor gene sets, indicating that ADM features broad acquisition of progenitor gene expression (Fig. S1K). Overall, inducible genes in ADM were divided evenly amongst those reflecting dedifferentiation to a late progenitor phenotype and those characteristic of a ductal transition (Fig. S2F-H). In this way, ‘ADM’ is not solely a ductal transition, but emerges from an unbiased developmental perspective as a broad acinar cell dedifferentiation to a late progenitor state.

### Enhancer reprogramming is induced by injury but not mutant Kras^G12D^

To understand epigenetic reprogramming elicited by injury, we FACS-sorted cells and performed the Assay for Transposase Accessible Chromatin with high-throughput sequencing (ATAC-seq) on early progenitor, late progenitor, acinar, and ductal cells, as well as acinar-derived ADM cells.

PCA analysis revealed that, from a chromatin standpoint, ADM again represents a hybrid ductal and late progenitor state (Fig. 1A). Interrogating all differentially accessible regions (DARs) between normal acinar and ADM cells, we observed strong commonalities of the latter with both late progenitor and ductal samples (P1/P2 and D1 clusters, respectively), while also retaining, to a lesser degree, some acinar features (A1/A2 clusters) (Fig. 1B). Among other benchmark cell types, ADM cells had the fewest DARs compared to acinar cells (n=19251) and late progenitors (n=46975), indicative of their dedifferentiated chromatin state (Fig. 1C). Examining the shared peaks between contrasts, we observed that ADM features robust acquisition of 5590 peaks found in both late progenitor and ductal cell types (Fig. 1D). Examining the content of the chromatin changes induced by injury, we observed strong enrichment of AP-1 motifs in gains of accessibility (Fig. 1E) and, using the Genomic Regions Enrichment of Annotations Tool (GREAT), observed Gene Ontology (GO) terms corresponding to MAPK signaling (Fig. 1F).

**Figure 1.**
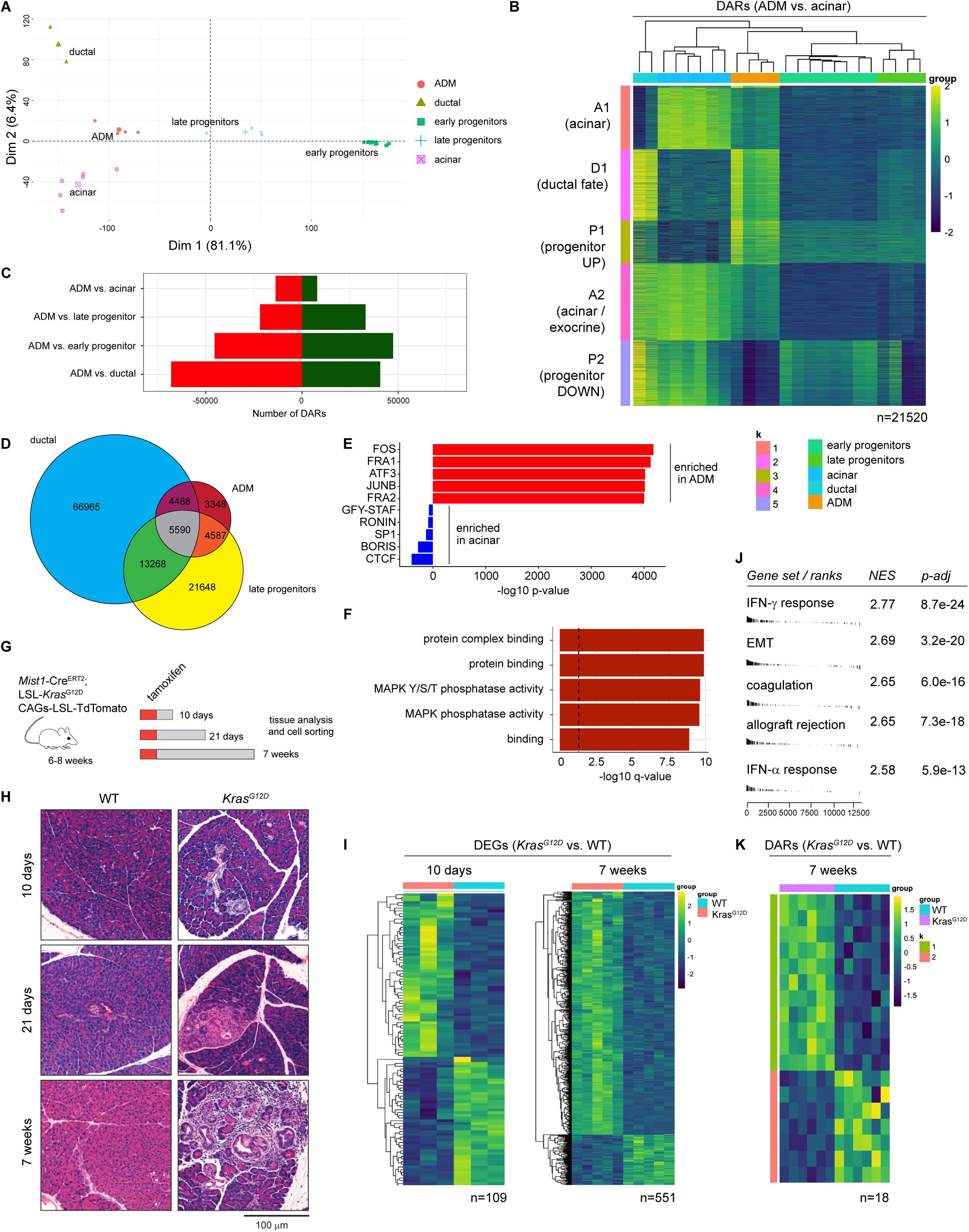
Injury, not mutant Kras, can initiate enhancer reprogramming in the acinar compartment. (A) PCA of ATAC-seq data for the top 5000 most variable genes in FACS-sorted early progenitor, late progenitor, acinar, ductal, and ADM cells. (B) Heatmap of differentially accessible regions (DARs; N = 21,520) identified after comparing ADM cells versus normal acinar cells, with chromatin features of ADM cells shown alongside late progenitor, ductal, and acinar cells. Thresholds: absolute log2(fold change) > 0; padj < 0.01. Each column is a biological replicate mouse. (C) Barplot quantifying the number of DARs between ADM and various benchmark cell types (acinar, ductal, early and late progenitors). Thresholds: absolute log2(fold change) > 0; padj < 0.01. (D) Venn diagram showing the overlap of DARs identified by comparing tdTomato+ cells isolated from WT acinar cells to each of the other cell types in the diagram (ADM, late progenitor, and ductal). Thresholds: absolute log2(fold change) > 1; padj < 0.01. (E) HOMER analysis depicting top motif enrichments when comparing ADM versus normal acinar. (F) Gene Ontology (GO) analysis using the Genomic Regions Enrichment of Annotations Tool (GREAT) on the regions with increased chromatin accessibility in ADM cells. (G) Schematic representation of the lineage-traced MK^G12D^T mouse model. (H) Hematoxylin and eosin staining of MK^G12D^T mouse pancreas sections after 10 days, 21 days and 7 weeks of mutant Kras activation. Representative images of N=3-5 mice per condition. Scale bar, 100 µm. (I) Heatmap of DEGs identified after comparing tdTomato+ cells isolated from WT to tdTomato+ cells collected from MK^G12D^T mice 10 days (N = 109) and 7 weeks (N = 551) of mutant Kras activation. Thresholds: absolute log2(fold change) > 0; padj < 0.01. Each column is a biological replicate mouse. (J) GSEA of RNA-seq data comparing normal acinar cells to tdTomato+ cells collected from MK^G12D^T mice 7 weeks of mutant Kras activation. (K) Heatmap of differentially accessible regions (DARs; N = 18) identified after comparing from MK^G12D^T mice 7 weeks of mutant Kras activation versus normal acinar cells. Thresholds: absolute log2(fold change) > 0; padj < 0.01. Each column is a biological replicate mouse.

Prior studies suggest that mutant *Kras* cooperates with injury to drive chromatin alteration in the pre-neoplastic pancreas.^5,11^ These studies, however, have leveraged embryonic *Kras* activation wherein robust progression to neoplasia is observed with *Kras* alone and superimposed inflammation is dispensable.^23–24^ In adult mice, mutant *Kras* alone is weakly tumorigenic, requiring prolonged periods of activation to form only sporadic neoplastic lesions, and caerulein for robust tumorigenesis.^8,23,25–26^ To examine whether mutant *Kras* can rewire chromatin in the adult mouse pancreas, we used MT; LSL-Kras^G12D^ (MK^G12D^T) mice to activate mutant *Kras*^G12D^ in differentiated acinar cells (Fig. 1G). In keeping with prior reports^26–27^, we observed only sporadic acquisition of ADM and PanIN up to 7 weeks after mutant *Kras* activation (Fig. 1H). We then performed RNA-sequencing of sorted tdT^+^ cells at 10 days and 7 weeks, finding that *Kras*^G12D^ expression results in 109 and 551 differentially expressed genes, respectively (Fig. 1I). GSEA revealed induction of Hallmark IFN-γ, IFN-α, and allograft rejection pathways, as well as EMT (Fig. 1J), with GO terms corresponding to immune and inflammatory responses (Fig. S3A). Despite these alterations to gene expression, there were only 18 DARs after 7 weeks of mutant *Kras* (Fig. 1K), suggesting that *Kras*-induced transcriptional responses are driven by mechanisms that do not involve direct changes in chromatin accessibility, such as the upregulation of pre-existing genes, activation of existing transcription factors, or post-translational modifications. Importantly, we did not find previously described *Kras*-induced DARs as dynamic in our model (Fig. S3B). As mutant *Kras* alone is not sufficient to trigger robust epigenetic remodeling, what has been described as a cooperativity with injury emerges instead as a dependency on injury-induced changes to chromatin.

### Kras^G12V^ and Kras^G12R^ are weak drivers of histologic transformation in the pancreas

In pancreatic cancer, G12D mutation correlates with poorer survival, while the G12R mutation is associated with better outcomes^19–20,28–29^, and early data suggest that G12V is also favorable.^19–20^ Embryonic *Kras*^G12R^ activation *in vivo* has been shown to be deficient as a potent inducer of premalignant disease^17^, but it is unclear how *Kras*^G12V^ and *Kras*^G12R^ compare to the well-studied *Kras*^G12D^ allele in tumor initiation. To systematically interrogate how the three most common PDAC-associated *Kras* variants drive tumorigenesis, we generated MK^G12Vgeo^T and MK^G12R^T mice (Fig. 2A). At 10 days, 21 days, and 7 weeks following mutant *Kras* activation, only *Kras*^G12D^ led to the development of sporadic ADM and PanIN (Fig. 2B-C). At 4 months, *Kras*^G12D^ progressed to exhibit more advanced and more numerous neoplastic lesions, which remained far fewer in *Kras*^G12V^ and *Kras*^G12R^ (Fig. 2D-F).

**Figure 2.**
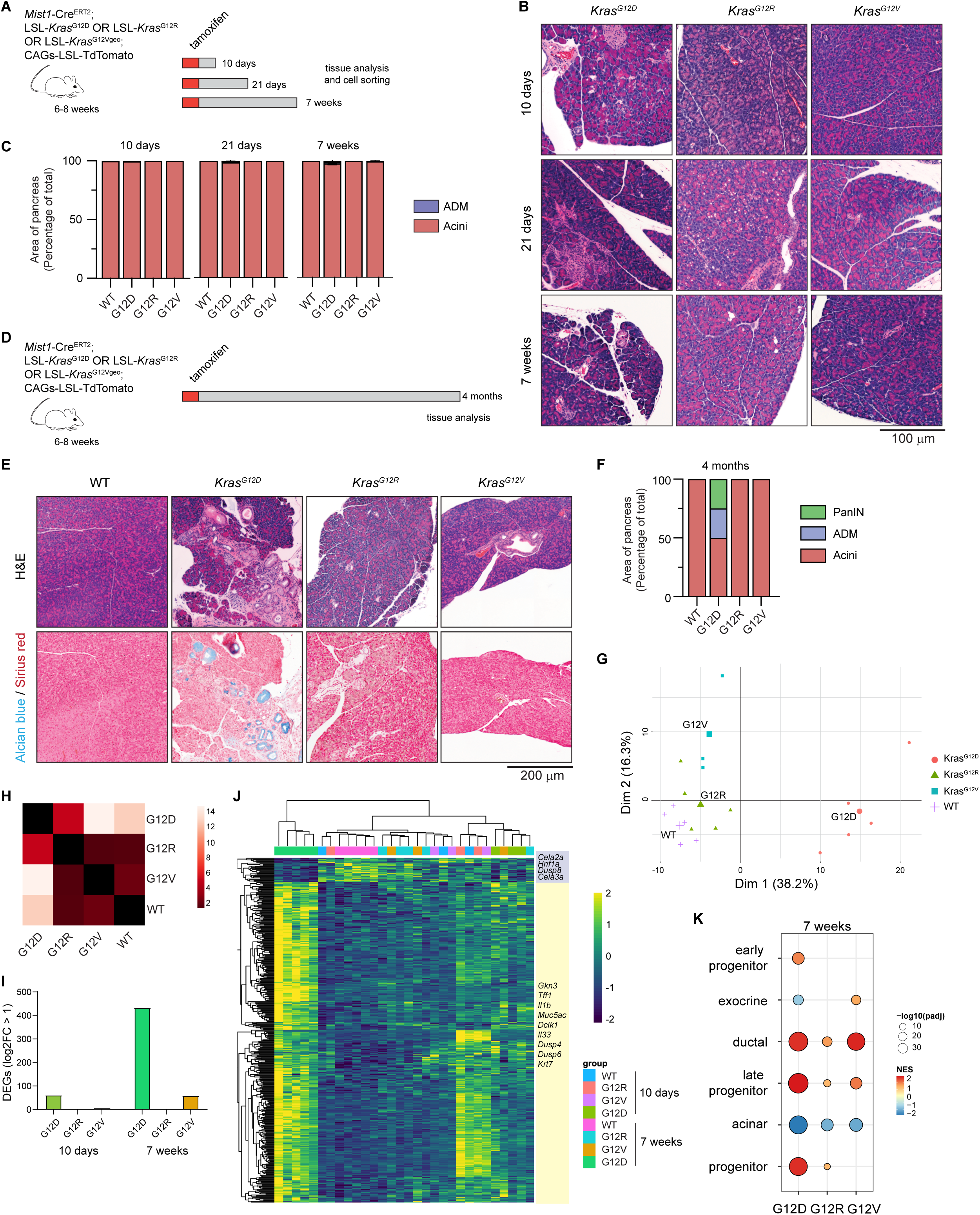
*Kras^G12V^* and *Kras^G12R^* are weak drivers of histologic transformation in the pancreas. (A) Schematic representation of the lineage-traced MK^G12D^T, MK^G12Vgeo^T, and MK^G12R^T mouse models after 10 days, 21 days, and 7 weeks of mutant Kras activation. (B) Hematoxylin and eosin (H&E) staining of MK^G12D^T, MK^G12Vgeo^T, and MK^G12R^T mouse pancreas sections collected after 10 days, 21 days, and 7 weeks of mutant Kras activation. Representative images of 2-5 mice per condition. Scale bar, 100 µm. (C) Histologic quantification of normal acinar tissue and ADM lesions in pancreata collected from MK^G12D^T, MK^G12Vgeo^T, and MK^G12R^T mice after 10 days, 21 days, and 7 weeks of mutant Kras activation. (D) Schematic representation of the lineage-traced MT (WT), MK^G12D^T, MK^G12Vgeo^T, and MK^G12R^T mouse models after 4 months of mutant Kras activation. (E) H&E and Alcian blue/Sirius red staining of pancreas sections collected from MT (WT), MK^G12D^T, MK^G12Vgeo^T, and MK^G12R^T mice after 4 months of mutant Kras activation. Images are representative of n = 1–6 mice per condition. Scale bar, 200 μm. (F) Histologic quantification of normal acinar tissue, as well as the percent area of ADM and PanIN lesions in pancreata collected from MT (WT), MK^G12D^T, MK^G12Vgeo^T, and MK^G12R^T mice after 4 months of mutant Kras activation. Data are summarized as mean ± SEM. Two-way ANOVA was performed. (G) PCA of RNA-seq data for the top 2000 most variable genes in tdTomato+ cells isolated from WT and tdTomato+ cells isolated from MK^G12D^T, MK^G12Vgeo^T, and MK^G12R^T mice after 7 weeks of mutant Kras activation. (H) Bhattacharyya distance analysis of the top 2000 most variable genes in tdTomato+ cells isolated from MT (WT), MK^G12D^T, MK^G12Vgeo^T, and MK^G12R^T mice after 7 weeks of mutant Kras activation. (I) Barplot quantifying the number of DEGs in MK^G12D^T, MK^G12Vgeo^T, and MK^G12R^T mice after 10 days and 7 weeks of mutant Kras activation. Thresholds: log2(fold change) > 1; padj < 0.01. (J) Heatmap of DEGs identified after comparing tdTomato+ cells isolated from WT to tdTomato+ cells collected from MK^G12D^T mice after 7 weeks (N = 551) of mutant Kras activation––these samples were visualized alongside MK^G12Vgeo^T and MK^G12R^T tdTomato+ cells collected from the same timepoint. Thresholds: absolute log2(fold change) > 0; padj < 0.01. Each column is a biological replicate mouse. (K) GSEA performed on RNA-seq data using adult and developing pancreas benchmark gene sets, independently comparing normal acinar cells to tdTomato+ cells collected from MK^G12D^T, MK^G12Vgeo^T, and MK^G12R^T mice after 7 weeks.

Across these models, we isolated tdTomato-positive cells at 10 days and 7 weeks and performed RNA-seq. PCA showed minimal separation at 10 days (Fig. S4A), and clear separation of *Kras*^G12D^ mutant acinar-derived cells from the other alleles by 6 weeks (Fig. 2G), which was also present at 7 weeks in Bhattacharyya distances for the top 2000 most variable genes (Fig. 2H). Interestingly, at 10 days, *Kras*^G12R^ cells were most distinct from *Kras*^WT^ (Fig. S4B), and gene pattern analysis of the 551 *Kras*^G12D^-induced DEGs at 7 weeks (Fig. 2I-J) showed that approximately 60% of these (Groups 2 and 4) were actually more highly expressed at the group level at the 10 day timepoint in *Kras*^G12R^ (Fig. S4C). By 7 weeks, *Kras*^G12R^ was not able to maintain any DEGs as compared to *Kras*^WT^ (Fig. 2I). Similarly, GSEA showed that *Kras*^G12R^ induced the most significant acquisition of progenitor gene expression at 10 days (Fig. S4D), but this was lost by 7 weeks, with only *Kras*^G12D^ robustly inducing acquisition of ductal and late progenitor transcriptional modules and loss of acinar identity programs (Fig. 2K). These findings suggest that *Kras*^G12R^ can initiate but not sustain mutant *Kras* signaling, with *Kras*^G12V^ as an intermediate potency mutant.

### Lineage plasticity is preserved in Kras^G12V^ and Kras^G12R^ mutants following injury

Given the requirement for inflammation to induce robust tumor initiation, we next asked how injury facilitates lineage plasticity across the *Kras* mutants. We induced mutant *Kras* and pancreatitis using tamoxifen and caerulein, respectively and interrogated mice at the peak of injury (2 days) and 21 days later (Fig. 3A). At the peak of injury (2d), all caerulein-treated *Kras* mutant mice exhibited robust formation of ADM (Fig. 3B) and induction of Sox9 and CK19 expression in metaplastic lesions (Fig. S5A), to a degree greater than *Kras*^WT^ and indistinguishable between the three *Kras* mutants (Fig. 3C).

**Figure 3.**
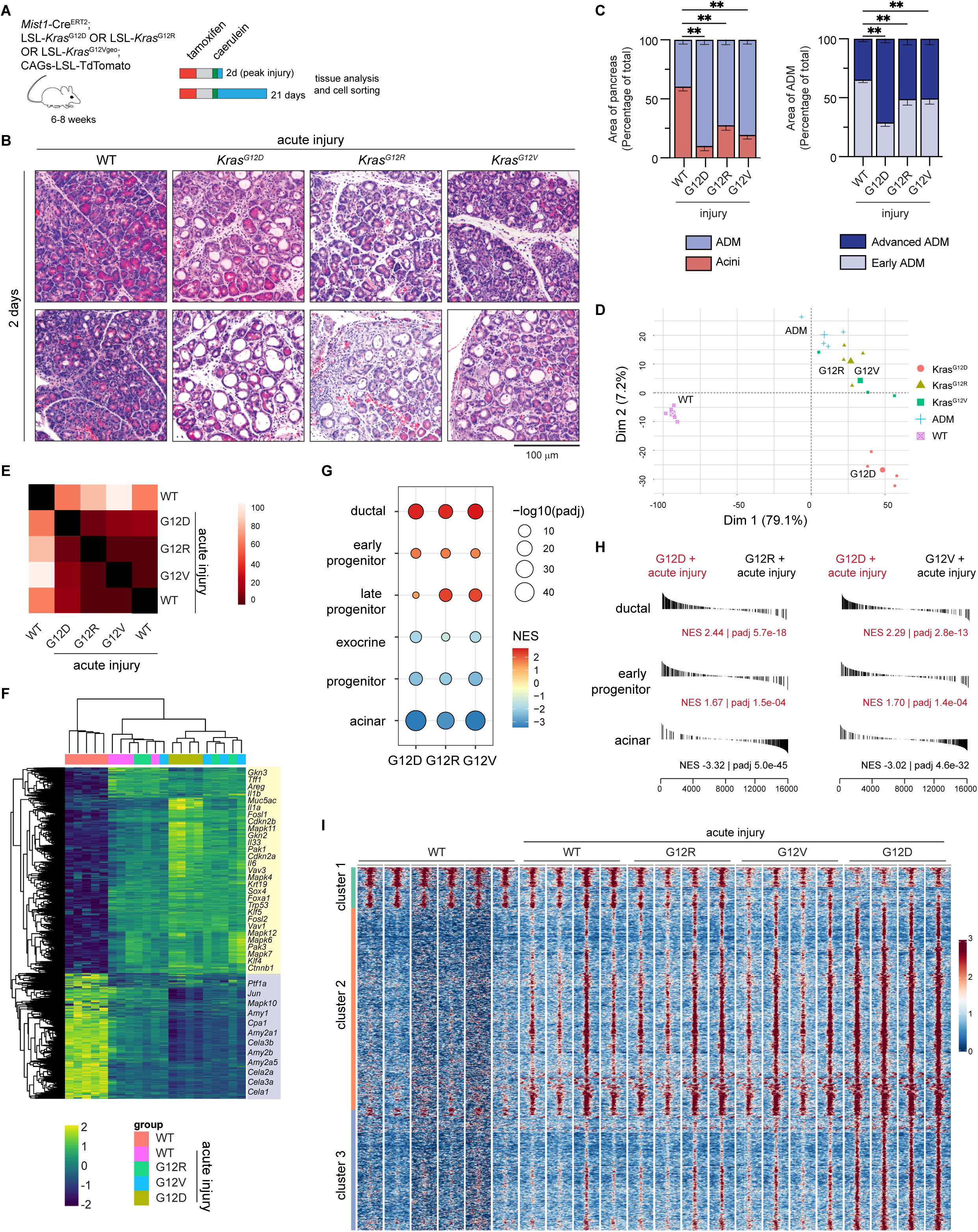
Lineage plasticity is not diminished in *Kras^G12V^* and *Kras^G12R^* mutants. (A) Schematic representation of the lineage-traced MT, MK^G12D^T, MK^G12Vgeo^T, and MK^G12R^T mouse models with acute injury following by 2 days and 21 days of recovery. (B) Hematoxylin and eosin staining of MT (WT), MK^G12D^T, MK^G12Vgeo^T, and MK^G12R^T mouse pancreas sections collected 2 days post-acute injury. Representative images of n=4-5 mice per condition. Scale bar, 100 µm. (C) Histologic quantifications of normal acinar tissue and ADM lesions in pancreata collected from MT (WT), MK^G12D^T, MK^G12Vgeo^T, and MK^G12R^T mice 2 days post-acute injury. ADM lesions further subdivided based on severity (early and advanced). (D) PCA of RNA-seq data for the top 2000 most variable genes in tdTomato+ cells isolated from WT without injury and tdTomato+ cells isolated from MT (ADM), MK^G12D^T, MK^G12Vgeo^T, and MK^G12R^T mice collected 2 days post-acute injury. (E) Bhattacharyya distance analysis of the top 2000 most variable genes in tdTomato+ cells isolated from MT (WT), MK^G12D^T, MK^G12Vgeo^T, and MK^G12R^T mice 2 days post-acute injury as well as from MT without injury. (F) Heatmap of all DEGs identified after comparing tdTomato+ cells isolated from WT without injury to tdTomato+ cells collected from acutely injured MT (WT), MK^G12D^T, MK^G12Vgeo^T, and MK^G12R^T mice 2 days post-injury. Thresholds: absolute log2(fold change) > 1; padj < 0.01. Each column is a biological replicate mouse. (G) GSEA performed on RNA-seq data using adult and developing pancreas benchmark gene sets, independently comparing acutely injured to tdTomato+ cells to MK^G12D^T-, MK^G12Vgeo^T-, and MK^G12R^T-derived tdTomato+ cells 2 days post-acute injury. (H) GSEA performed on RNA-seq data using adult and developing pancreas benchmark gene sets, independently comparing tdTomato+ cells collected from MK^G12D^T mice to both MK^G12Vgeo^T and MK^G12R^T mice 2 days post-acute injury. (I) Heatmap of all differentially accessible regions identified after comparing control MT cells (WT) with MK^G12D^T cells 2 days post-acute injury. Thresholds: absolute log2(fold change) > 0; padj < 0.01. Each column is a biological replicate mouse.

To define how peak injury and oncogenic Kras synergize to drive lineage plasticity, we isolated lineage-traced acinar-derived cells in WT, *Kras*^G12D^, *Kras*^G12V^, and *Kras*^G12R^ contexts, 2 days after the completion of caerulein. PCA of RNA-sequencing showed mild separation of *Kras*^G12D^ populations (Fig. 3D) from other mutant *Kras* alleles only in Dim2 (7.2% of variance), and Bhattacharyya distances between the three mutants were quite small (Fig. 3E). Similarly, *Kras* mutants were highly similar across the ∼6000 DEGs, with the most dramatic differences in gene expression present between all injury contexts (whether wild-type or mutant for *Kras*) and control acinar cells (Fig. 3F). However, comparison of the different *Kras* mutants to each other showed that each was able to upregulate ductal transcripts but that G12D was able to do so to a greater degree, and could drive re-expression of the early progenitor module and to repress acinar cell identity, with G12V and G12R retaining the late progenitor features associated with ADM (Fig. 3G-H). Gene pattern analysis showed that the vast majority of DEGs were either induced or repressed in the context of ADM and the degree of change was mildly enhanced in the context of *Kras*^G12R^, more markedly altered in *Kras*^G12V^, and most significantly changed in *Kras*^G12D^ (Fig. S5B), defining a hierarchy amongst the three alleles.

To understand how *Kras*^G12D^ and *Kras*^G12R^ diverge molecularly at the peak of injury despite similar histology, we carefully interrogated differences in gene expression between these two alleles. Along with several MAPK pathway genes (*Mapk3* [ERK1]*, Mapk11, Mapk6, Map2k1* [MEK1], we observed that *Klf5*, implicated in the progenitor program induced by injury^5^, and IL-33, a nuclear cytokine that supports tumorigenesis^11^, induced in G12D, are not upregulated in G12R (Fig. S5C). Conversely, *Bhlha15* (Mist1), a key acinar identity TF was retained in G12R (Fig. S5C). Pathway analysis showed enrichment of KRAS signaling, TGF-β, and EMT gene sets in G12D as compared to both G12V and G12R (Fig. S5D). Finally, enriched GO terms in G12D corresponded to actin filament / cytoskeletal organization processes (Fig. S5E), findings that lined up with enrichment of the apical junction Hallmark in G12D (Fig. S5D), suggesting a diminished capacity for G12R to alter epithelial polarity and the actin cytoskeleton in tumor formation.

Next, we examined how the different *Kras* alleles are able to rewire chromatin in the context of caerulein. Chromatin accessibility profiling largely recapitulated the gene expression findings, with *Kras*^G12D^ most markedly driving gains of accessibility amongst gains shared with *Kras*^G12R^ and *Kras*^G12V^ (Fig. 3I; cluster 2); interestingly, *Kras*^G12D^ was also associated with an allele-specific cluster of gains in ATAC signal (Fig. 3I; cluster 3). Analysis of these clusters revealed enrichment of ATF3 and AP-1 motifs in chromatin domains unveiled specifically in the context of *Kras*^G12D^ (Fig. S5F). Together these data highlight the unique ability of *Kras*^G12D^ to drive tumorigenic epigenetic programs and malignancy-associated transcriptional programs. Indeed, when we examined mice 21 days after injury, the histology of *Kras*^G12D^ diverged from the other *Kras* alleles, with *Kras*^G12V^ giving rise to occasional PanIN (as opposed to more widespread PanIN for *Kras*^G12D^). *Kras*^G12R^ at this time point was largely indistinguishable from WT mice that were examined 21 days after caerulein (Fig. S5G-H), reflecting the ultimate phenotypic consequence of the molecular differences observed at 2 days.

### Kras^G12D^ is unique in its ability to drive robust neoplastic transformation

As initial lineage plasticity was comparable between *Kras*^G12D^, *Kras*^G12V^, and *Kras*^G12R^ immediately following acute caerulein, we asked if prolonged subacute caerulein-mediated injury could sustain neoplastic transformation in *Kras*^G12V^ and *Kras*^G12R^. To this end, we administered caerulein using a 3-week protocol, and then harvested the pancreas at either peak injury (2 days), 21 days, or after a recovery interval of 3 months (Fig. 4A). Surprisingly, prolonged inflammation could not support *Kras*^G12V^ or *Kras*^G12R^ tumorigenesis across all timepoints (Fig. 4B), with a striking separation of G12D from the other alleles that was more enhanced over time (Fig. 4C). In survival analyses, G12D mice uniformly did not survive beyond 24 weeks (owing not to tumor but to pancreatic insufficiency), whereas WT, G12V, and G12R mice generally survived >1 year (Fig. 4D).

**Figure 4.**
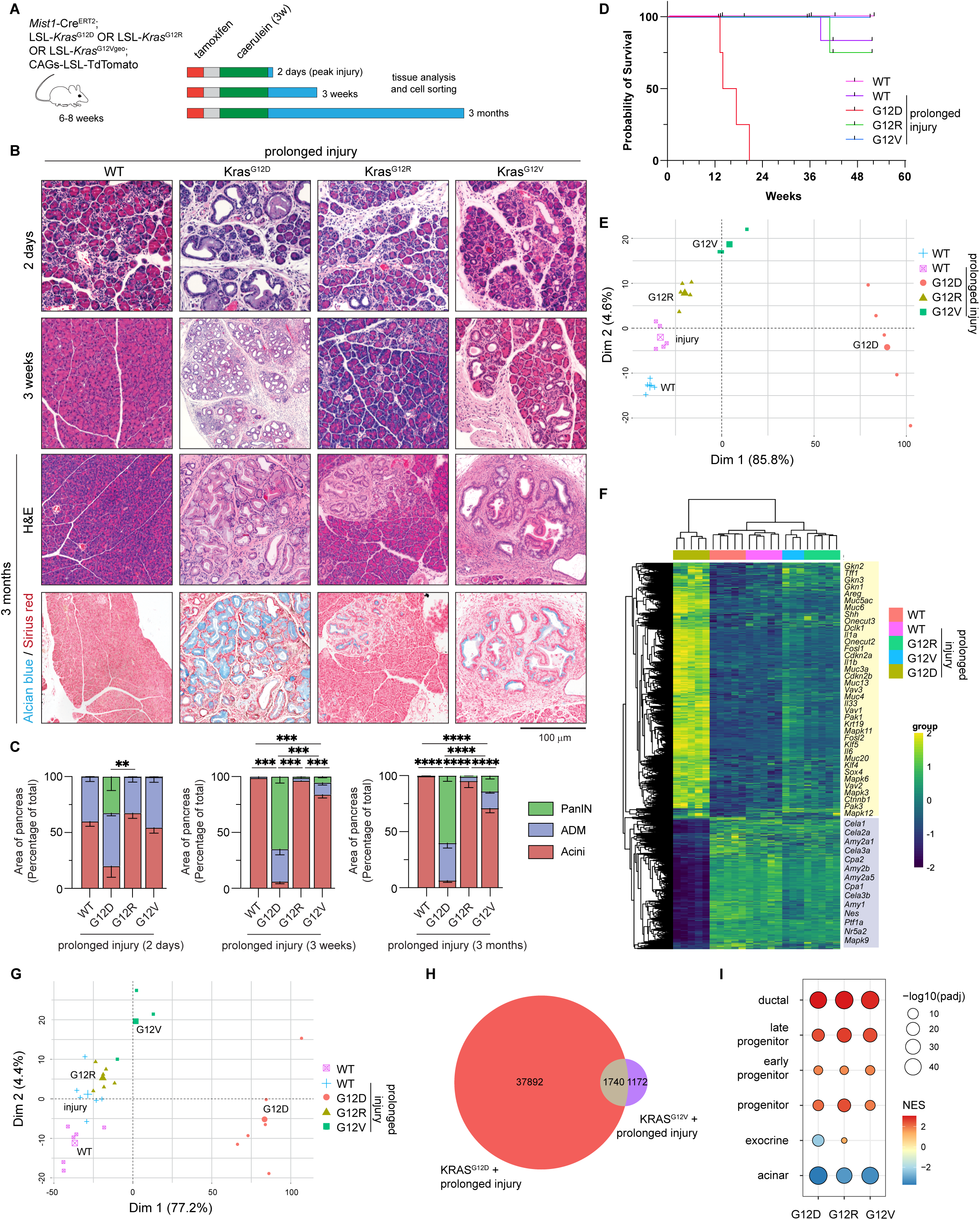
*Kras^G12D^* is unique in its ability to drive robust neoplastic transformation. (A) Schematic representation of the lineage-traced MT, MK^G12D^T, MK^G12Vgeo^T, and MK^G12R^T mouse models after prolonged injury, as well as 2 days, 3 weeks, and 3 months of recovery. (B) Hematoxylin and eosin staining of MT, MK^G12D^T, MK^G12Vgeo^T, and MK^G12R^T mouse pancreas sections collected 2 days, 3 weeks, and 3 months post-prolonged injury. Alcian blue/Sirius red staining is also shown for the 3 months timepoint. Representative images of n=3-5 mice per condition. Scale bar, 100 µm. (C) Histologic quantifications of normal acinar tissue, ADM, and PanIN lesions in pancreata collected from MT, MK^G12D^T, MK^G12Vgeo^T, and MK^G12R^T mice 2 days, 3 weeks, and 3 months post-prolonged injury. (D) Kaplan-Meier curves of overall survival for MT, MK^G12D^T, MK^G12Vgeo^T, and MK^G12R^T mice after prolonged injury. Add number of mice for each genotype. (E) PCA of RNA-seq data for the top 2000 most variable genes in tdTomato+ cells isolated from WT without injury and tdTomato+ cells isolated from MT, MK^G12D^T, MK^G12Vgeo^T, and MK^G12R^T mice collected 3 weeks post-prolonged injury. (F) Heatmap of all DEGs identified after comparing tdTomato+ cells isolated from WT without injury to tdTomato+ cells collected from MK^G12D^T mice 3 weeks post-prolonged injury. Thresholds: absolute log2(fold change) > 1; padj < 0.01. Each column is a biological replicate mouse. (G) PCA of ATAC-seq data for the top 2000 most variable regions in tdTomato+ cells isolated from WT without injury and tdTomato+ cells isolated from MT, MK^G12D^T, MK^G12Vgeo^T, and MK^G12R^T mice collected 3 weeks post-prolonged injury. (H) Venn diagram showing the overlap of DARs identified by comparing tdTomato+ cells isolated from both MK^G12D^T and MK^G12Vgeo^T collected 3 weeks post-prolonged injury. Thresholds: absolute log2(fold change) > 1; padj < 0.01. (I) GSEA performed on RNA-seq data using adult and developing pancreas benchmark gene sets, independently comparing tdTomato+ cells isolated from MT mice without injury to tdTomato+ cells collected from MK^G12D^T, MK^G12Vgeo^T, and MK^G12R^T mice 3 weeks post-prolonged injury.

As before, we sorted acinar-derived cells from WT, *Kras*^G12D^, *Kras*^G12V^, and *Kras*^G12R^ mice, all treated with prolonged caerulein, 3 weeks after the completion of injury. In this context, G12D transcriptionally was quite different from the other alleles, both in PCA (Fig. 4E) and when interrogating DEGs in the set (Fig. 4F). When evaluating the ATAC-seq data, PCA again showed that the G12D mutant separates away from the other Kras mutant samples and control samples (Fig. 4G), with 39,632 DARs as compared to 2,912 for G12V and 0 for G12R (Fig. 4H).

When we examined the transcriptional programs driving this divergence between alleles, we observe that at the pathway level, all three mutant *Kras* alleles could drive expression of ductal and progenitor gene modules (Fig. 4I), suggesting that lineage reversion and transdifferentiation programs were uniformly inducible. Similarly, *Kras*^G12R^ and *Kras*^G12V^ were able to drive EMT and inflammatory signaling (IL-6 signaling, inflammatory response, complement pathway) Hallmarks to a similar degree as *Kras*^G12D^ (Fig. S6A). Notably, however, *Kras*^G12R^ and *Kras*^G12V^ were deficient in their ability to repress exocrine / acinar cell fate (Fig. 4I). Direct comparison of *Kras*^G12D^ to the other mutants revealed that G12D drove TNF signaling via NF-κB and apical junction Hallmarks to a much greater degree, with the latter not at all enriched in G12R (Fig. S6B). GO term analysis on DEGs supported the distinct ability once again of G12D to drive transcriptional programs related to cytoskeletal / actin filament organization (Fig. S6C). In keeping with these findings, we observed in the ATAC data that G12D and to a lesser extent G12V, but not G12R, were able to induce PDAC-associated epigenetic reprogramming (Fig. S6D). Together, these data define a phenotypic defect in G12R, and to a lesser extent in G12V, not to change identity but to sustain specific oncogenic transcriptional and epigenetic programs as robustly as G12D.

### Kras^G12R^ and Kras^G12V^ feature impaired EGFR signaling

As our data supported the presence of a clear hierarchy amongst the PDAC-associated *Kras* variants, we sought to understand why the ‘extremes’ in our analysis - *Kras*^G12R^ and *Kras*^G12D^ - drive markedly different molecular and phenotypic outcomes. Using the Cytokine Signaling Analyzer (CytoSig)^30^, we parsed our gene expression data to define G12D-specific cytokine signaling cascades, observing that EGF and TGF-α family signaling are strongly induced in G12D but not as well in the other allelic contexts (Fig. S7A). Examination of the most enriched cytokine transcripts in the context of G12D nominated amphiregulin (AREG) and epiregulin (EREG) as key EGFR ligands that are G12D-specific in acute injury and more highly enriched as phenotypes diverge in the context of prolonged injury (Fig. S7B). To confirm that these ligands are in fact more highly present in G12D at the protein level, we performed a multiplexed cytokine array that revealed a 50-fold induction of AREG in G12D, compared to ∼2-fold in G12R and G12V (Fig. S7C). Importantly, several key cytokines elaborated in PDAC initiation were also mildly enriched in the G12D setting, including IL-6, LIF, and Cxcl1/2 (Fig. S7D).

To ascertain if the difference in *Kras* mutants could be ‘rescued’ by EGFR ligand administration, we turned to a well-established model of *in vitro* ADM^1,13^ using recombinant human TGFα administered to acinar explants. We observed more robust ductal cyst formation in *Kras*^G12D^ acinar cells than in either *Kras*^G12V^ or *Kras*^G12R^ (Fig. S7E-F), highlighting that the differences among *Kras* alleles are not driven solely by a paucity in EGFR ligand. Mutant KRAS^G12D^ is known to drive increased expression of EGFR itself^31^, and, indeed, neoplastic transformation is known to depend on the presence of signal through the receptor.^31–32^ To determine if G12V and G12R can also drive EGFR upregulation, we performed Western blotting, finding that both EGFR and phosphorylation at Y1068 were both induced in G12D but not in G12V/R (Fig. S7G-H) though the proportion of phospho-EGFR relative to total appeared unchanged across the mutants (Fig. S7I). Collectively, these data support a multi-level defect in G12V and G12R to drive both EGF ligand and receptor that are necessary for KRAS-mediated oncogenic transformation.

### Activation of Rac1 signaling is deficient in the Kras^G12R^ mutant pancreas

Next, we evaluated the signaling implications of dampened EGF receptor-ligand interactions at the plasma membrane. Because *Kras*^G12R^ has been described to have different GTP hydrolysis activity^33–34^, we first examined RAS-GTP levels in the mouse pancreas 7 weeks after tamoxifen injection (as in Fig. 3A). All three mutant alleles increased active RAS to comparable degrees above wild-type, with minimal difference between them (Fig. 5A; Fig. S8A). We next examined MAPK signaling leading to ERK activation. Here we observed in the adult acinar-specific system ERK phosphorylation tended to be elevated *in vivo* in the context of G12D in the absence of injury (Fig. 5B; Fig. S8B) and more significantly in acute injury (Fig. S8C) as compared to G12R. However, in the context of prolonged injury – timepoints at which there is phenotypic divergence – there were similar levels of p-ERK between G12D and G12R, with G12V actually showing the most Erk activation (Fig. S8D). Indeed, by IHC, G12R showed a capacity to generate p-ERK^+^ PanIN, albeit with significant temporal delay (Fig. 5C), suggesting that MAPK signaling is delayed but not deficient in G12R when neoplastic lesions do form.

**Figure 5.**
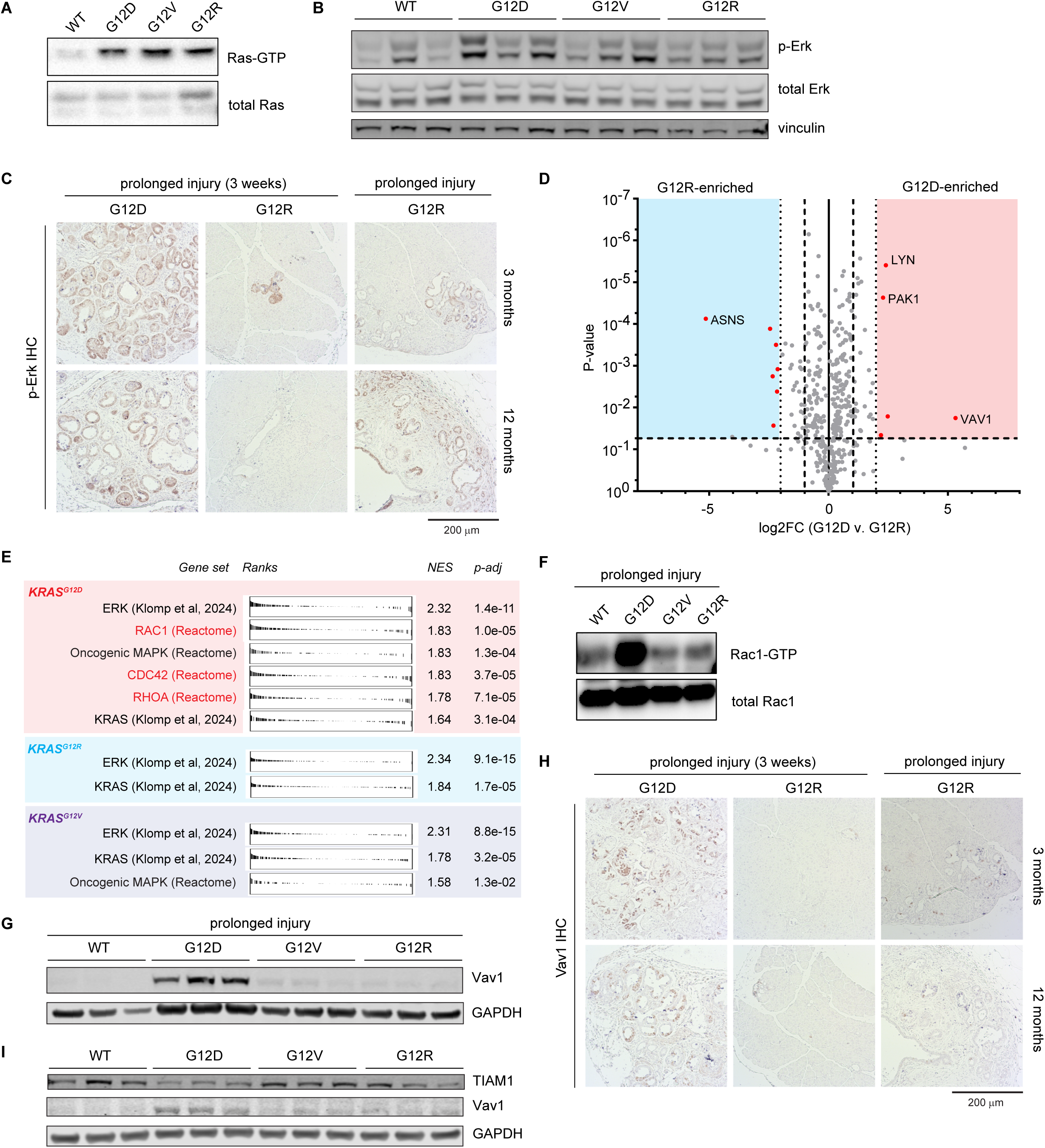
Activation of Rac1 signaling is deficient in the *Kras*^G12R^ mutant pancreas. (A) Ras-GTP levels from bulk pancreas tissue lysate extracted from MT, MK^G12D^T, MK^G12Vgeo^T, and MK^G12R^T mice 7 weeks post-tamoxifen injection. (B) Western blot showing Akt phosphorylation, total Akt and β-actin of bulk pancreas tissue lysate extracted from MT, MK^G12D^T, MK^G12Vgeo^T, and MK^G12R^T mice 7 weeks post-tamoxifen injection. (C) Levels of Erk phosphorylation, total Erk and vinculin from bulk pancreas tissue lysate extracted from MT, MK^G12D^T, MK^G12Vgeo^T, and MK^G12R^T mice 7 weeks post-tamoxifen injection. (D) Immunohistochemical (IHC) staining of pancreatic sections for phospho-Erk in both MK^G12D^T and MK^G12R^T mice 3 weeks, 3 months, and 12 months after prolonged injury. Representative images of n=2-5 mice per condition. (E) Reverse-phase protein array (RPPA) analysis identified VAV1 and PAK1 as differentially abundant proteins between MK^G12D^T and MK^G12R^T 3 weeks after prolonged injury, performed on n=3 mouse pancreas lysates per condition. (F) GSEA performed on RNA-seq data comparing tdTomato+ cells isolated from WT without injury to tdTomato+ cells collected from MK^G12D^T (top), and MK^G12R^T (middle), MK^G12Vgeo^T (bottom) mice 3 weeks post-acute injury. (G) Rac1-GTP levels from bulk pancreas tissue lysate extracted from MT, MK^G12D^T, MK^G12Vgeo^T, and MK^G12R^T mice 3 weeks after prolonged injury. (H) Western blot showing Vav1 and GAPDH expression levels from lysates obtained from sorted acinar-derived (tdTomato+) cells extracted from MT, MK^G12D^T, MK^G12Vgeo^T, and MK^G12R^T mice 3 weeks after prolonged injury. (I) IHC staining of pancreatic sections for VAV1 in both MK^G12D^T and MK^G12R^T mice 3 weeks, 3 months, and 12 months after prolonged injury. Representative images of n=2-5 mice per condition. (J) Western blot showing TIAM1, VAV1 and GAPDH expression levels from sorted tdTomato+ cells collected from MT, MK^G12D^T, MK^G12Vgeo^T, and MK^G12R^T mice 7 weeks post-tamoxifen injection.

To ascertain in an unbiased fashion the upstream signaling differences between *Kras*^G12D^ and *Kras*^G12R^, we performed a reverse-phase protein array (RPPA) on bulk pancreas lysates obtained at 3 weeks following prolonged injury (as in Fig. 4A). Much to our surprise, we observed that VAV1 and PAK1 were enriched in G12D (Fig. 5D), at ∼40-fold and ∼6-fold, respectively, more than G12R, along with asparagine synthetase (ASNS), enriched in G12R. VAV1 is well-described as a key guanine nucleotide exchange factor (GEF) for the Rho GTPase RAC1^35^, which in turn activates PAK1 and p38/JNK. Our GSEA and GO analyses suggested a G12R loss of apical junction and actin remodeling (Fig. S6B-C), processes in which RAC1 signaling is essential and that are required for PDAC initiation.^36–37^ Indeed, we observed in the RNA-sequencing data from mice that gene sets related to Rac1 signaling pathways, but not gene sets relating to MAPK/ERK signaling nor the recently developed KRAS molecular signature^38^, were enriched in G12D relative to G12R and G12V (Fig. 5E). Given the well-established role for RAC1 activity in key cytoskeletal processes and PAK activation, our data pointed to the RAC1 pathway as different between G12D and G12R.

Next, we directly queried if active RAC1 was different between *Kras* alleles. In bulk pancreas lysates, using a GST-PAK1-protein binding domain(PBD), we detected much higher RAC1-GTP in G12D than in G12V or G12R (Fig. 5F). We then validated the observed difference between G12D and G12R in VAV1 abundance (found in the RPPA) by Western blotting (Fig. 5G; Fig. S8E). As *Vav1* is expressed highly in the hematopoietic compartment, we confirmed by IHC that the G12D pancreatic epithelium was VAV1^+^, as in prior reports^39^, and that this staining was confined largely to PanIN lesions. In addition, the G12R pancreas was largely devoid of VAV1 staining, and PanIN that eventually emerged failed to stain robustly for VAV1 (Fig. 5H).

Finally, we asked if VAV1 differences across mutant *Kras* alleles preceded phenotypic divergence between the alleles. Indeed, VAV1 abundance showed marked difference between G12D and G12R in the context of *Kras* activation alone (at 7 weeks) (Fig. 5I; Fig. S8F), a context in which there is limited histologic transformation. This difference was also observed in sorted tdTomato^+^ cells, indicating that the difference between G12D and G12R in VAV1 is established early in the acinar (and acinar-derived) compartment (Fig. S8G). However, in the context of acute injury, wherein the *Kras* alleles produce similar phenotypes (Fig. 3B), we observed no difference in VAV1 abundance (Fig. S8H), suggesting that VAV1 can initially be articulated in G12R in initial lineage plasticity, but cannot be maintained over time (Fig. 5H). Importantly, VAV1 has been shown to be dependent on EGFR^39^, connecting the inability to initiate the circuit of autocrine AREG upregulation and EGFR activation to diminished RAC1 signaling. Our data thus converge on the notion that VAV1 - previously described to be a key node in pancreatic tumorigenesis^39–41^, and also lost in the absence of p110α^40^ – is not activated in the context of *Kras*^G12R^, and that the failure of G12R to mediate oncogenic transformation is due to a lack of EGFR/VAV1 supporting sustained RAC1 activity that is required for enforcement of the neoplastic cell fate decision.

### Constitutive activation of AKT can support Kras^G12R^-driven tumorigenesis

In light of the inability of *Kras*^G12R^ to drive robust tumorigenesis, we aimed to determine if this inability could be overcome by secondary perturbations *in vivo*. We turned to the well-established ‘KPC’ model of PDAC featuring *p48-*Cre and mutant p53 (LSL-*Trp53*^R172H^) to evaluate if tumor suppressor loss could ‘rescue’ *Kras*^G12R^ (Fig. 6A). Surprisingly, at 20 weeks of age, the G12R mouse pancreas was devoid of lesions (Fig. 6B) at a time point when G12D mice already developed robust PanIN. Mouse survival was substantially prolonged (Fig. 6C), but beginning at approximately 40 weeks of age, several G12R/p53^mut^ mice developed lymphomas or required euthanasia for other conditions. In total, of the 11 G12R mice that died in the course of the study, 6 had evaluable pancreas histology, none of which contained PanIN lesions or PDAC. In contrast, all G12D mice did not survive >40 weeks. We suspected that biallelic inactivation of p53 might potentiate neoplastic commitment, and generated MK^G12R^T; LSL-*Trp53*^R172H/flox^ mouse models accordingly (Fig. 6D). Strikingly, whereas G12D;p53^mut/flox^ animals rapidly developed PanIN and PDAC by 20 weeks of age (Fig. 6E), G12R;p53^mut/flox^ animals were free of neoplastic lesions and showed robust overall survival (Fig. 6F).

**Figure 6.**
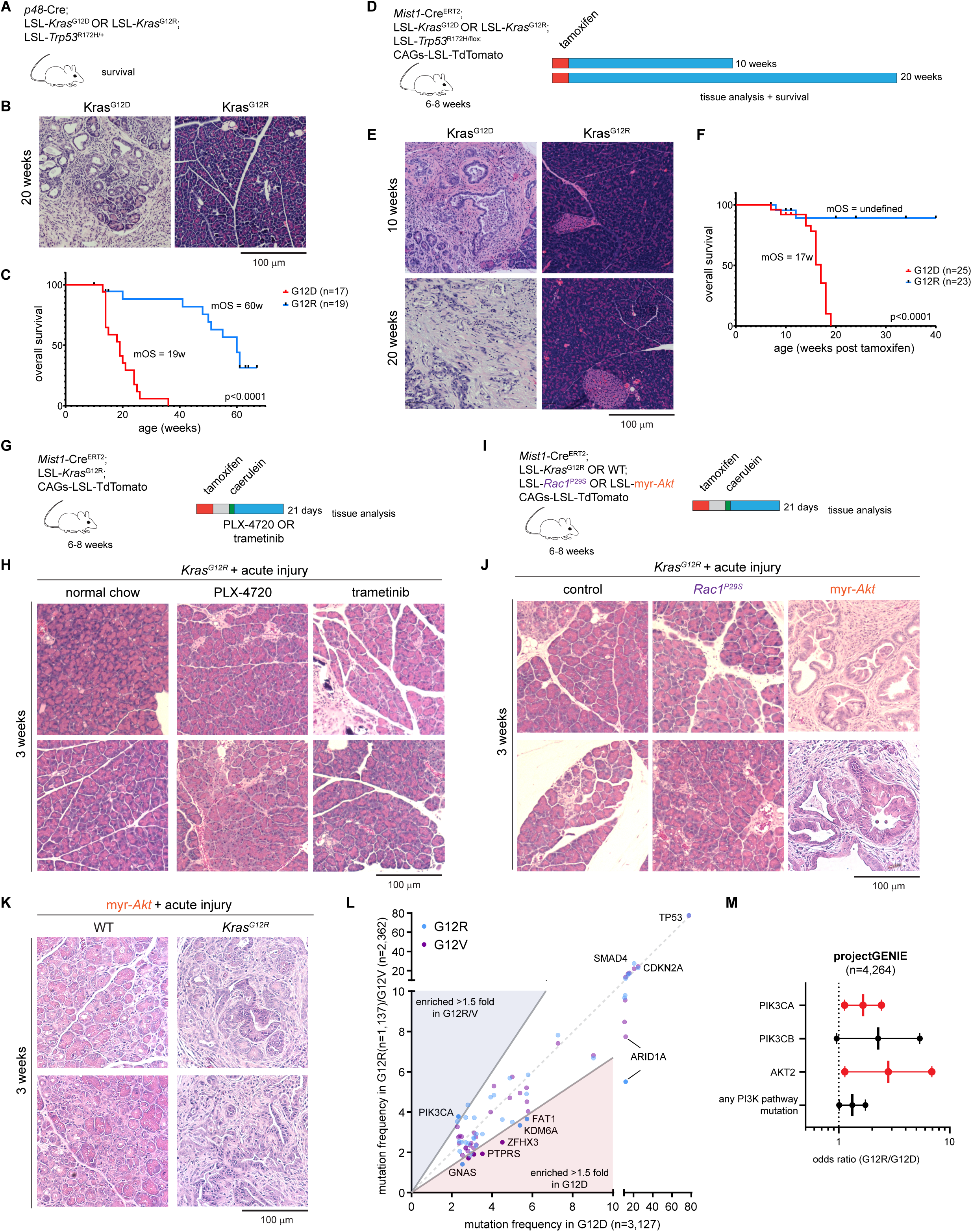
Constitutive activation of AKT can support Kras^G12R^-driven tumorigenesis. (A) Schematic representation of p48-Cre; LSL-*Kras*^G12D/R^; LSL-*Trp53*^R172H/+^ mouse model. (B) Representative hematoxylin/eosin sections of mouse pancreas from animals as described in (A). (C) Kaplan-Meier survival of mice as per (A); Mantel-Cox test was performed. (D) Schematic representation of the lineage-traced MK^G12R^T and MK^G12D^Tmouse models with *Trp53*^R172H/flox^ alleles. (E) Hematoxylin/eosin staining of pancreas from animals as described in (A), at the indicated timepoints. (F) Kaplan-Meier survival of mice as per (D); Mantel-Cox test was performed. (G) Schematic representation of the lineage-traced MK^G12R^T mouse models after acute injury +/-low-dose trametinib or PLX-4720 diet for 3 weeks, with evaluation at 3 weeks of recovery from injury. (H) Hematoxylin and eosin staining of MK^G12R^T mouse pancreas sections collected 3 weeks post-acute injury and the indicated treatment. (I) Schematic representation of the lineage-traced MK^G12R^T +/- *Rac1^P29S^* OR *myr-Akt* mouse models after acute injury, with evaluation at 3 weeks of recovery from injury. (J) Hematoxylin and eosin staining of MK^G12R^T +/- *Rac1^P29S^* OR *myr-Akt* mouse pancreas sections collected 3 weeks post-acute injury. (K) Hematoxylin and eosin staining of MT;*myr-Akt* +/- *Kras*^G12R^ mouse pancreas sections collected 3 weeks post-acute injury. (L) Scatter plot of the mutational frequency (%) of the indicated genes in *KRAS*^G12D^ (x-axis) and *KRAS*^G12R^ / *KRAS*^G12V^ tumors (y-axis), using the projectGENIE dataset (www.cbioportal.org). (M) Forest plot indicating the relative odds ratio (G12R/G12D) for co-occurring mutation in in the indicated gene, using the projectGENIE dataset.

Given the heterogeneity among KRAS mutants to interact with RAF kinases, we next asked if modulation of MAPK activation could ‘rescue’ the G12R defect. To test this, we perturbed MAPK activation in MK^G12R^T mice through either administration of low-dose MEK inhibitor (trametinib; 0.5mg/kg) or RAF inhibitor (PLX4720; 472 mg/kg) to either diminish or paradoxically amplify^42–44^ ERK signaling, respectively, addressing the possibility that *Kras*^G12R^ might lead to too much or too little signaling through this output (Fig. 6G). Tissue analysis after 3 weeks of the addition of trametinib or PLX-4720 to the diet, however, showed no significant PanIN development in G12R mice (Fig. 6H).

We turned our attention to the key signaling defects in G12R nominated previously and herein. Prior data has highlighted that *Kras*^G12R^ interacts more poorly with the PI3K subunit p110α^18^ with deficient AKT activation with ectopic expression in both mouse^17^ and human^18^ fibroblasts; however, a separate analysis of the mouse pancreas following embryonic *Kras* activation showed no difference in levels of phospho-AKT.^17^ Despite this, as PI3K can also directly activate RAC1^40^, we suspected that either constitutive activation of AKT or RAC1 could ‘rescue’ the ability to drive PanIN in the *Kras*^G12R^ context. To test this directly, we crossed a myristoylated AKT (myrAKT) allele^45^ or the constitutively active *Rac1*^P29S^ allele^46^ with MK^G12R^T mice (Fig. 6I). While constitutive RAC1 activation supported tumor initiation, we found that enforcement of membrane localization of AKT could indeed rescue the G12R phenotype (Fig. 6J). As activating PI3K mutations have been shown to independently drive pancreatic tumorigenesis, even in the absence of mutant *Kras*^47^, we confirmed that while myrAKT alone gave rise to ADM and scattered PanIN, the addition of KRAS^G12R^ dramatically potentiated neoplastic commitment (Fig. 6K). These data show that constitutive activation of AKT can cooperate with KRAS^G12R^ to drive tumorigenesis.

In light of these findings, we suspected that human *KRAS*^G12R^ would display evidence of deficient oncogenic signaling and a requirement for PI3K/AKT/RAC1 activation. We queried the projectGENIE cohort of >8,000 PDAC patients to determine if G12V/G12R mutation co-occurs with other mutational or copy number alterations. While most mutational events, including to *TP53, SMAD4,* and *CDKN2A*, did not display a predilection for a specific *KRAS* allele (Fig. 6L), *PIK3CA* mutations were enriched in *KRAS*^G12R^ (Fig. 6L-M). Indeed, we observed a trend towards statistically significant enrichment of all PI3K pathway gene mutations, including *PIK3CB* and *AKT2* (Fig. 6M). These data support the notion that in human disease there are defects in KRAS^G12R^ signaling correlating with the mouse models that show an inability to support RAC1 activity, its key GEF VAV1, and in AKT activation, all of which are downstream of the KRAS-PI3K interaction (Fig. 7).

**Figure 7.**
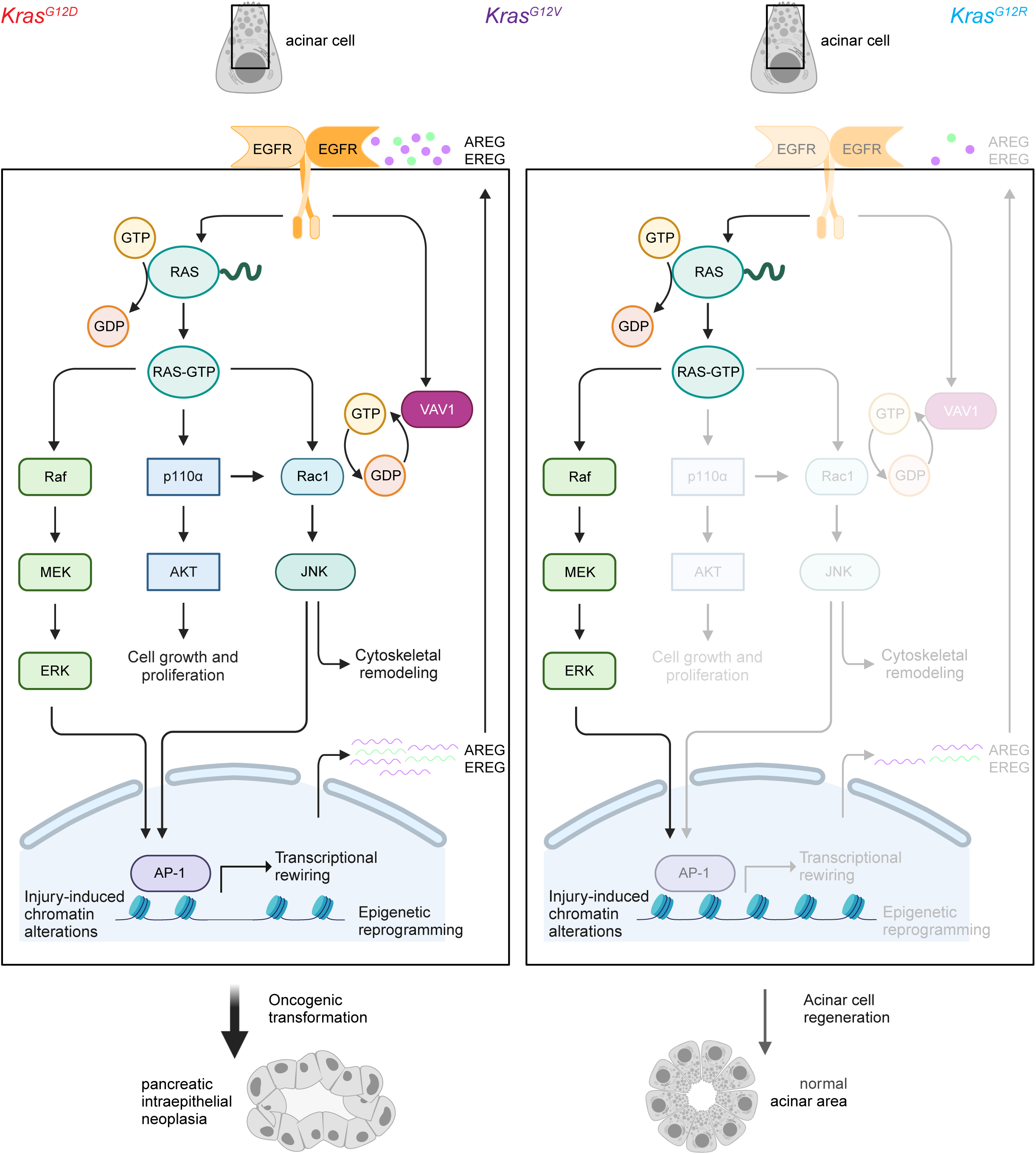
An emergent hierarchy among *Kras* mutant alleles in the pre-neoplastic pancreas. Schematic summary of the findings of the current study. Created with Biorender.

## DISCUSSION

Cancer initiation is a complex process wherein mutational events in association with exogeneous insults can synergistically overcome biological barriers that maintain tissue homeostasis. In the context of pancreatic cancer, the driver genetic events have long been understood, such that these have been deployed for 20 years to establish *in vivo* models that recapitulate the phenotypic behavior of human disease and uncover multiple molecular dependencies of pancreatic cancer initiation and progression.

Despite this, there have been several limitations of the robust scientific evidence arising from the most frequently utilized models. For the most part, Cre drivers have largely consisted of *Pdx1-*Cre or *p48-*Cre, which induce recombination events in the embryonic pancreas. Tumorigenesis in these contexts does not require superimposed inflammation. Adult models of mutant *Kras* activation (such as the *Ptf1a-*Cre^ER^ model) do show a role for exogenous injury, but still generate a haploinsufficient state in *Ptf1a*, which has been shown to destabilize acinar cell identity.^48^ Lastly, there has been widespread use of lineage-tracer alleles (such as mTmG and rtTA-IRES-mKate [RIK]) whose effects on homeostasis in the pancreas have been recently called into question.^17^ Physiologic models that feature adult *Kras* activation, show a requirement for injury, and that preserve differentiated cell states – such as is feasible in the *Mist1-*Cre^ERT2^ system employed herein – have not been deployed broadly across the field.

In addition, across several models, injury has been suggested to not just provoke ADM, but the accumulation of varying subsets of acinar cells characterized by different markers. These include TFF2^49^, DCLK1^50^, Nestin^1^, Stmn1^51^, TERT^52^, and KLF5^5^, the latter being suggestive of a progenitor program unveiled in the acinar lineage. To date, however, this work has largely been marker-specific, such that there has not been a broad interrogation of the acinar-progenitor dedifferentiation that occurs in injury. A third limitation of the extant literature has been the near-exclusive focus on *Kras*^G12D^. KRAS^G12V^ and KRAS^G12R^ – representing nearly half of all PDAC cases – have been far less well-studied, and have recently in our work emerged as prognostic of patient outcome and biologically distinct in the human setting.^19^

In our work, we have attempted to comprehensively address these limitations. Specifically, we have interrogated the acinar cell fate in injury and tumor initiation, with a view to (1) contextualizing the injury response against originating acinar fate, ductal cells, and the early and late progenitor compartments; (2) understanding the specific transcriptional and epigenetic effects of *Kras* and/or injury in a physiologic model; and (3) articulating the specific effects of each of the 3 most common *Kras* mutations, G12D, G12R, and G12V. Through a combination of lineage-tracing models and molecular analyses, and reconstruction of the developmental trajectory of exocrine cells of the pancreas, our work has revealed that injury-induced ADM reflects a mixed-lineage state that can be viewed as representing both a ductal metaplasia and a reversion to a late progenitor phenotype. This state, arising in the injury response to caerulein, features broad epigenetic reprogramming; mutant *Kras*, by contrast, induces oncogenic signaling and alterations to gene expression, but does not lead to any substantial rewiring of chromatin. Through this lens, we find that mutant *Kras* and injury do not so much as cooperate as it is that *Kras* co-opts the lineage reversion brought about by injury.

This co-optation of lineage reversion is robust for *Kras*^G12D^ with induction of profound epigenetic reprogramming, characterized by an accumulation of accessibility in AP-1 chromatin and a repression of acinar cell identity. In contrast, *Kras*^G12V^ and *Kras*^G12R^ cannot leverage the mixed-lineage state induced by injury to drive robust tumorigenesis. At the pathway level, *Kras*^G12R^ instead can induce an initial degree of gene expression alteration that can support lineage plasticity, but does not sustain either a histologic transformation, transcriptional rewiring, nor epigenetic reprogramming.

There are a multitude of mechanisms by which *Kras*^G12D^, *Kras*^G12V^, and *Kras*^G12R^ mutants could give rise to varied phenotypes. Biochemical analyses have shown that G12D and G12V comparably lower the rate of GTP hydrolysis, and that G12R displays the most dramatic reduction as compared to WT KRAS^34^, suggesting that KRAS^G12R^ might be more locked into its active conformation. However, the KRAS mutants also vary in their GEF interactions, with SOS1 catalytic constants (*k*_cat_) lowest for G12R and GRP1 *k*_cat_ lowest for G12D.^18^ As SOS1 is the dominant GEF for KRAS, these data imply that G12R might instead be less ‘active’ than G12D by virtue of an inability to engage in rapid GTP-GDP exchange. In our data, we observe that RAS-GTP levels appear largely comparable between the *Kras* mutants (but elevated above wild-type) to where these biochemical differences do not appear to translate into a broad difference in active RAS.

In addition to changes to GTP hydrolysis and exchange, KRAS mutants have also been described to differ in their interaction with downstream effectors. RAF affinity has been described as decreased in all three KRAS variants as compared to WT.^34^ In contrast, KRAS^G12R^ leads to a switch II domain structural change, such that this mutant appears to interact poorly with p110α.^18^ In transformed fibroblasts and human PDAC cell lines, this defective interaction of KRAS^G12R^ is associated with a lack of PI3K-Akt-mTOR signaling. Additionally, it has been suggested that other PI3K isoforms can substitute for the absence of p110α activation. Direct evidence for this has been provided by prior work with pancreas-specific *Kras*^G12D^ p110α-knockout mice, wherein a similar phenotypic outcome of an inability to sustain injury-induced metaplasia was seen; in this context, there was a defect in VAV1 upregulation and sustained RAC1 signaling^40^, closely recapitulating the findings in our G12R model. At first blush, our data suggest that the differences in phenotypic and molecular outcomes between the *Kras* mutants can be explained by differences in RAC1 / VAV1 activation, pathways that are integral to cell migration and cytoskeletal dynamics, are dependent on EGF receptor-ligand interactions, and that are enriched in G12D but not the other alleles. Yet despite this, we find that constitutive activation of RAC1 signaling via the P29S mutant does not ‘rescue’ the G12R phenotype, but enforced AKT signaling does. These data thus imply that the G12R defect is multifactorial, consisting of both inadequate proliferative signaling through AKT as well as in RAC1 activation.

A key implication of our data extends to therapeutic targeting of *KRAS* in each allelic context. Our data demonstrate that mutant *KRAS* supports the pleiotropic downstream effectors to varying degrees. As multiple lines of evidence support roles for MAPK, PI3K, and RAC1 pathway activation in tumor initiation, one would suspect that G12D, capable of supporting all three signaling outputs might be more dependent on mutant KRAS, while G12R in particular might be less susceptible to KRAS targeting. Some evidence supporting this notion emerge from recent evaluation of the RMC-6236 pan-RAS(ON) inhibitor, where PDX/CDX models showed 0% (0/4) disease control rate (DCR) for G12R models but 56% (5/9) for G12D.^53^ Indeed, whether there proves to be mutant-specific variation in the response to KRAS targeting remains an open question.

Taken together, our work herein underscores the complex cell fate decisions that typify the responses to inflammatory injury and oncogenic stress. Whereas these two essential components conspire to drive oncogenesis - even with broad temporal separation^13^ – they can be viewed as serving unique, non-overlapping roles in this process. Injury induces the requisite epigenetic reprogramming that *Kras* cannot; *Kras*, in turn, can co-opt the mixed-lineage state to sustain a stable neoplastic cell fate decision. Moreover, whereas lineage plasticity is facilitated by all three *Kras* mutants, neoplastic commitment depends significantly on the precise mutation in *Kras*, suggesting that tumorigenesis in each mutational context has distinct allele-specific molecular dependencies, and perhaps therapeutic vulnerabilities.^18^ Quite clearly, *KRAS* allelic status can shape clinical outcomes in human pancreatic cancer by driving distinct biological behavior; here we demonstrate that the basis for differences in cancer begin with distinct evolutionary trajectories preceding tumorigenesis. Moving forward, these differences highlight the diverse biological implications of *Kras* mutations – across preclinical models, human disease, and pharmacologic targeting of KRAS -- that warrant further study to enable the rational and precise interception of pancreatic cancer.

## ACKNOWLEDGMENTS

This work was supported by an American Association for Cancer Research-Pancreatic Cancer Action Network Pathway to Leadership Award (RC), NIH/NIDDK R01 DK060694 (AKR/RC), NIH/NCI R01 CA204228 (RC/SDL), and NIH/NCI R01 CA290616 (RC/LED). AG is supported by the Prevent Cancer Foundation. This research was also funded in part through the NIH/NCI Cancer Center Support Grant P30 CA008748. Generation of the Ptf1a-tdT mouse was supported by NIH/NIDDK R01 DK020593 (to Alvin Powers, Vanderbilt University) and NIH/NIDDK R01 DK072473 (MAM).

## AUTHOR CONTRIBUTIONS

*Conceptualization:* AG, DJF, SDL, AKR, LED, RC; *Methodology:* AG, DJF, CWC, GP, MC, WJS, AO, YM, MPZ, WBF, AFR, MAM, SDL, AKR, LED, RC; *Software*: DJF, PZ, TMY, AFR, DB; *Validation*: AKR, LED; *Formal analysis:* AG, DJF, PZ, TMY, AFR, EH, RKY, DB; *Investigation:* AG, DJF, CWC, GP, MC, WJS, AO, YM, MPZ, WBF, LED, RC; *Resources*: MC, MPZ, MAM, LED, RC; *Data curation:* AG, DJF, LED, RC; *Writing – original draft*: AG, DJF, RC; *Writing – review & editing*: AG, DJF, LED, RC; *Visualization*: AG, DJF, RC; *Project administration*: DB, MAM, SDL, AKR, LED, RC; *Supervision:* DB, MAM, SDL, AKR, LED, RC; *Funding acquisition:* AG, DJF, MAM, SDL, AKR, LED, RC.

## METHODS

### Animal models

Mice were housed in a pathogen-free facility at Weill Cornell Medicine (WCM) under standard housing conditions. Mouse lines used were described previously: Mist1-Cre^ERT2^ (MGI:3821734 Bhlha15^tm3(cre/ERT2)Skz^)^24^, LSL-Kras^G12D^ (MGI:2429948 Kras^tm4Tyj^)^54–55^, LSL-Kras^G12R^ (MGI:6281560 Kras^em2Ldow^)^17^, LSL-Kras^G12V^ (MGI:3582830 Kras^tm1Bbd^)^56^, R26^LSL-tdTomato^ (MGI:3809523 Gt(ROSA)26Sor^tm9(CAG-tdTomato)Hze^)^57^, p48-Cre (MGI:2387812 Ptf1a^tm1.1(cre)Cvw^)^58^, LSL-Trp53^R172H^ (MGI:3039264 Trp53^tm2.1Tyj^)^59^, LSL-Trp53^fl^ (MGI:193101; Trp53^tm1Brn^)^60^. Mice were bred to generate Mist1-Cre^ERT2^, LSL-Kras^G12D or G12R or G12V^, R26^LSL-tdTomato^ referred to as MK^G12D^T, MK^G12R^T, MK^G12V^T mice, with or without p53^R172H^ or p53^flox^ alleles, as indicated. Mice without any of the Kras mutants (G12D, G12R and G12V) are referred to as MT mice. Male and female animals were used for experiments. Ptf1a-Tdt mice (MGI:6470589 Ptf1a^tm3.1Mgn/Vu^), generated in this study are deposited at the Vanderbilt Cryopreserved Mouse Repository (https://vcmr.vcscb.org/mouse.html?vger_id=MB). Rac1^P29S^ mice^46^ were obtained from The Jackson Laboratory (RRID:IMSR_JAX:033790); myrAkt mice^45^ were generously provided by T. Wunderlich (Max Planck Institute for Metabolism Research). All manipulations were performed under the Institutional Animal Care and Use Committee (IACUC)–approved protocol (2017-0038) at Weill Cornell Medicine.

### Primary acinar cell culture

Acinar 3D cultures were generated as described^61^ with few modifications. Briefly, acinar cell isolation was obtained with collagenase/dispase mix dissociation as described below, then cells were filtered through a 100 μm cell strainer.

### Generation of Ptf1a-Tdt mice

Ptf1a-tdT mice independently expresses tdTomato along with an N-terminal tandem Strep-tag II /Flag tagged version of Ptf1a from the endogenous locus. ES cells containing the Ptf1a Loxed Cassette Acceptor (Ptf1aLCA) allele were used. This allele contains a LoxP site and an inverted LoxP site inside the Ptf1a locus. These sites allow for recombinase mediated cassette exchange. The exchange vector replaced exon 1 of Ptf1a with tdTomato, in frame with a viral 2A peptide, followed by an N-terminal tandem Streptag II/Flag tag fused to the Pft1a cDNA sequence. The Ptf1a^LCA^ allele was used to insert the Ptf1a^tdTomato-2A-NSF-ptf1(+HygroR)^ cassette using recombinase mediated cassette exchange. Following confirmation of exchange via sequencing, the mice were crossed to a flippase (FLP) recombinase carrying mouse to remove the Hygromycin resistance gene. tdTomato expression is evident in adult acinar cells, but is absent from islets, ducts, and the vasculature. The pancreas appears entirely normal by H&E and mice survive as homozygotes, implying an entirely functional Ptf1a protein. This novel mouse strain facilitates any experimentation requiring visualization of embryonic pancreatic progenitors cells or adult acinar cells, as well as immunoprecipitation of Ptf1a protein using the FLAG tag. For genotyping, the wildtype allele has a 636 bp band, while the transgenic allele has a 670 bp band. This allows us to distinguish wild-types, heterozygotes and homozygotes. Forward primer: CCTTCTGACTTCTCCAAGAAGGCA; Reverse primer: CCCTTTATGCCTGGCATTTCACTG.

### Tamoxifen treatment

Tamoxifen (Sigma T2859-5G) was dissolved in corn oil and administered by subcutaneous injections (at the indicated ages) at a dosage of 5 mg per injection. Mice were injected once a day for a total of 3 days––administered every other day (total duration: 5 days). Mice were allowed to recover 1 week after the last Tamoxifen treatment before receiving other treatments (if outlined in schema).

### Experimental pancreatitis

For acute injury treatment, mice received 8 hourly intraperitoneal (IP) injections over two consecutive days of either PBS (saline) or caerulein at a dosage of 75 μg/kg diluted in sterile PBS. For the prolonged injury treatment, mice were treated with three hourly IP injections a day, three times a week (on alternating days) for three weeks. For each experiment, either whole pancreata, or a portion of pancreata, was harvested for histologic analysis and downstream -omics assays.

### Embryonic pancreatic dissociation

Wildtype C57BL/6 and Ptf1a-tdT mice were used in all experiments. The fertilization of pregnant females was time-stamped with the discovery of vaginal plugs; Day 0 of embryonic life was assumed to have taken place at noon on the day vaginal plugs were discovered (i.e., days post coitum). Pregnant females were euthanized at the appropriate gestational age with embryos being removed from the uteri, and then placed in a Petri dish containing PBS. Embryos were carefully removed from the uteri with the assistance of a dissection microscope. At E10.5, the excised tissue included pancreatic rudiments (tdT-positive), as well as gut and stomach (tdT-negative). While at E15.5, the pancreas was excised free of surrounding gut and stomach. The excised tissue from both E10.5 and E15.5 were processed into a single-cell suspension using 1 mL of Accumax (Invitrogen, Cat: 00-4666-56) along with approximately 4 µL of 5 mg/mL DNase I [0.02 mg/mL] to minimize cellular clumping. Each tube contained a maximum of 10 embryos to avoid inefficient dissociation. The samples were then incubated in a 37°C water bath for 1 hour, during which the tubes were gently agitated by tapping and trituration to aid in dissociation. After incubation, the tubes were placed on ice and 1 mL of quenching solution was added to each sample (ingredients: supplemented L15 media [see below], DNase I [37.5 µg/mL]) was added to each sample. The tissues were then triturated 7-8 times with a P1000 filtered pipette to achieve thorough dissociation. The resulting cell suspension was filtered through a 35 µm mesh filter into FACS tubes, and the original tube was rinsed with double the volume of diluted quenching solution (ingredients: supplemented L15 media [see below], DNase I [7.5 µg/mL]) to maximize cell recovery. The filtered cell suspensions were centrifuged at 1100 rpm for 10-15 minutes at 4°C, and the supernatant was carefully aspirated, leaving 50-100 µL at the bottom of each tube. Cells were resuspended in quenching solution with DAPI [1:100] for approximately 15-30 minutes before FACS. All samples were processed for FACS analysis immediately after staining to ensure cell viability. Note: the supplemented L15 media used throughout was freshly prepared (ingredients: L15 media w/o phenol red (Gibco, Cat: 21083027), HEPES [10 mM], BSA [1 mg/mL], Pen/Strep [1%]). Similarly, both the concentrated and diluted quenching solutions were prepared fresh.

### Mouse pancreatic dissociation

Mice were euthanized using CO_2_, and pancreata were excised and placed in 3 mL of ice-cold HBSS as remaining pancreata were being harvested. Then, the ice-cold HBSS solution was discarded, and pancreata were placed in a 10 cm dish, containing 5 mL of 37°C collagenase and dispase (CD) solution––collagenase D is used for normal tissue; collagenase V is used for fibrotic tissue (CD ingredients: HBSS w/ Ca^2+^ Mg^2+^, Collagenase D / V [1 mg/mL], Dispase II [2 U/mL], STI [0.1 mg/mL], DNase I [0.1 mg/mL]). While in CD solution, pancreata were mechanically dissociated into ∼1-3 mm pieces using razor blades. Minced pancreata were then individually transferred to 50 mL conical tubes, and an additional 5 mL of CD solution was added, bringing each sample to a final volume of 10 mL. Tubes were placed on a 37°C orbital shaker at 135 rpm for 25 minutes. After incubation, the samples were centrifuged at 1000 rpm for 3 mins, washed with PBS, and then resuspended and incubated with 2 mL of 0.05% Trypsin-EDTA (37°C) for less than 3 min. Trypsin was inactivated immediately after incubation with 10 mL of 37°C PBS/FBS solution (ingredients: PBS w/o Ca2+ Mg2+, FBS [1:5], STI [0.1 mg/mL], DNase I [0.1 mg/mL]). Each tube was then gently inverted twice, centrifuged, and resuspended in 10 mL of 37°C FACS buffer (ingredients: PBS, EGTA [10mM], FBS [2%], STI [0.1 mg/mL], DNase I [0.1 mg/mL]). Cell suspension was passed through a 100 μm mesh filter into a new 50 mL conical tube, centrifuged, resuspended with 1 mL of FACS buffer solution with DAPI staining [1:100], and then, finally, transferred into a 40 μm FACS filter tube. All samples were kept on ice before and after FACS. For all FACS, Becton-Dickinson Influx or Becton-Dickinson Aria II were used to collect cells of interest. DAPI-/tdTomato+ cells were collected for MT, MK^G12D^T, MK^G12R^T, and MK^G12Vgeo^T mice. For ductal cell isolation, cells were stained prior to FACS with Cd49f and Cd133. DAPI-/Cd49f+/Cd133+ cells were collected as ductal cells while DAPI-/CD49f+ cells were collected as tdTomato-negative acinar cell controls.

### Bulk RNA-seq

Following mouse pancreas dissociation, DAPI-/tdTomato+ cells were FACS-sorted directly into TRIzol (100k cells per 750 mL TRIzol) in 1.5 mL microcentrifuge tube. Guanidinium thiocyanate-phenol-chloroform extraction was performed directly and the quality of the samples was determined using the Agilent RNA 6000 Nano Kit on the Agilent Bioanalyzer (Weill Cornell Medicine Genomics Resources Core Facility; WCM-GRCF). The WCM-GRCF prepared libraries using TruSeq RNA Sample Preparation (Non-Stranded and Poly-A selection), and used the NovaSeq 6000 (S1 Flow Cell – Paired End 2×50 cycles) for sequencing. The sequences were aligned to the mouse reference genome (mm9) using STAR4, a universal RNAseq aligner. To improve accuracy of the mapping, the genome was created with a splice junction database based on Gencode vM1 annotation.^62^ Sequences that mapped to more than one locus were excluded from downstream analysis, since they cannot be confidently assigned. Uniquely mapped sequences were intersected with composite gene models from Gencode vM1 basic annotation using featureCounts^63^, a tool for assigning sequence reads to genomic features. Composite gene models for each gene consisted of the union of exons of all transcript isoforms of that gene. Uniquely mapped reads that unambiguously overlapped with no more than one Gencode composite gene model were counted for that gene model; the remaining reads were discarded. The counts for each gene model correspond to gene expression values, and were used for subsequent analyses. Prior to the detection of differentially expressed genes, the quality of the sequences was assessed based on several metrics using FastQC^64^ and QoRTs^65^. Differential gene expression analysis was performed for each comparison using limma voom with default parameters.^66^

### Bulk ATAC-seq

Following pancreas cell isolation, 50k DAPI-/tdTomato+ acinar cells were sorted into FACS buffer supplemented with FBS [1:10].^67^ Briefly, acinar cells were centrifuged at 500g for 5 minutes (4°C) and washed with 50 μL of cold PBS. Cells were subsequently lysed using cold lysis buffer, and immediately spun at 500g for 10 min (4°C). The pellet was resuspended in the transposase reaction mix for the Tn5 tagmentation step for 30 minutes at 37°C and sample was purified using a Qiagen MinElute PCR Purification Kit. Next, DNA was indexed and amplified using PCR. The quality of the samples was assessed using Agilent High Sensitivity DNA kit. DNA libraries were then multiplexed and sequenced on a NextSeq2000 (Paired End; P2 - 100 cycles). Data analysis, including quality and adapter filtering, was applied to raw reads using ‘trim_galor’ before aligning to mouse assembly mm9 with bowtie2 using the default parameters. The Picard tool MarkDuplicates (http://broadinstitute.github.io/picard/) was used to remove reads with the same start site and orientation. The BEDTools suite (http://bedtools.readthedocs.io) was used to create read density profiles. Enriched regions were discovered using MACS2 and scored against matched input libraries (fold change > 2 and FDR-adjusted p-value < 0.1). A consensus peak atlas was then created by filtering out blacklisted regions (http://mitra.stanford.edu/kundaje/akundaje/release/blacklists/mm9-mouse/mm9-blacklist.bed.gz) and then merging all peaks within 500 bp. A raw count matrix was computed over this atlas using featureCounts (http://subread.sourceforge.net/) with the ‘-p’ option for fragment counting. The count matrix and all genome browser tracks were normalized to a sequencing depth of ten million mapped fragments. DESeq2 was used to classify differential peaks between two conditions using fold change > 2 and FDR-adjusted p-value < 0.1. Peak-gene associations were made using linear genomic distance to the nearest transcription start site with Homer (http://homer.ucsd.edu).

### Acinar 3D cell culture preparation

Following isolation of acinar cells, cell pellets were resuspended in acinar-explant (AE) media composed of DMEM supplemented with 1% FBS, 1% PS, 1% STI and 1mg/ml of Dexamethasone and then mixed with Matrigel (Corning) in a 2:1 ratio cells:Matrigel. Per 24-well plate, 400 μl of the cells:Matrigel suspension were plated and incubated at 37 °C for solidification for 2-3 hours. Upon Matrigel solidification, 400 μl of warm AE media was added with or without TGF-α at 50ng/ml concentration. Media (with and without TGF-α) was changed at day 1 and day 3 of culture. Brightfield image at 4X and 10X magnification were taken using a Nikon ECLIPSE Ti inverted microscope system equipped with an Andor Zyla 5.5 sCMOS camera.

### Analysis of brightfield images of spheroids with computer vision

Images were acquired in OME-TIFF format with 16-bit depth in grayscale. Illumination was normalized by subtracting the image passed by a gaussian filter of sigma = 5 * max(x_dim, y_dim) / 300, where x/y_dim are the image dimensions. Images were then inverted and intensity values clipped to the intensity between 3 and 99.8 percentiles, scaling image intensity values to unit range. To segment the spheroid objects from the background, we detected edges in the image by Sobel filtering, and seeded the watershed algorithm using values below 0.3 as background and above 0.95 as positive. We dilated objects in this binary mask with a disk of 3 pixels of diameter, and filtered objects smaller than 32 pixels in diameter. We removed further smaller objects by applying closing (dilation followed by erosion), filling holes within objects, and applying erosion and dilation again, all with a 3 pixel diameter disk. All operations were performed using scikit-image version 0.18.2.11 We then quantified various features for each object in each image using the ‘skimage.measure.regionprops’ function, and reduced values per image using the mean. Statistical testing was performed between groups of interest with a two-sided Mann–Whitney U test, and adjusted for multiple comparisons with the Benjamini–Hochberg False Discovery Rate method using pingouin (version 0.4.0).^68^

### Histopathology and immunofluorescence staining

Pancreata were fixed overnight in 4% buffered PFA, transferred to 70% ethanol, and then embedded in paraffin using IDEXX BioAnalytics laboratory. Serial sections were cut, and then hematoxylin and eosin (H&E) staining was performed. For IF, slides were deparaffinized, underwent an antigen retrieval using Sodium citrate buffer, blocked with 5% BSA supplemented with 0.4% Triton X-100 in PBS, and primary antibodies were incubated overnight at 4°C. Secondary antibodies conjugated with Alexa-488 or Alexa-647 (Invitrogen) were used and DAPI nuclear counterstaining was performed. Fluorescent images were captured with a Nikon ECLIPSE Ti inverted microscope system equipped with an Andor Zyla 5.5 sCMOS camera or entire slide/region area where digitalized with a Zeiss Axioscan 7. For Alcian blue staining, we used the Alcian Blue (ph2.5) Stain Kit (#H-3501) following the manufacturer’s recommendation. H&E and Alcian blue images were captured with a ZEISS Axio Scope.A1 equipped with a Axiocam 105 color or entire slide/region area where digitalized with a Zeiss Axioscan 7.

### Gene set enrichment analysis

Using normalized read counts of RNA-seq data, the fgseaMultilevel function from the fgsea package with the following parameters were used: minGSSize = 15; maxGSSize = 500 MSigDB Hallmark, MSigDB KEGG, MSigDB REACTOME, and MSigDB Biocarta gene sets were tested for all analyses. Genes were ranked based on DESeq’s wald statistic (stat), which takes into account the log-fold change and its standard error.

### Sample preparation and RPPA

Mice were euthanized using CO_2_, and pancreata were excised, washed quickly in 5 mL of ice-cold HBSS to remove excess blood, put into a 1.5 mL microtube and tube was submerged into liquid nitrogen for snap freezing. Briefly, a total of 23 samples, MT, MTK^G12D^, MTK^G12R^, MTK^G12V^ mice after saline or caerulein treatment 3 weeks after treatment were processed. Proteins in cellular lysates were denatured using a buffer solution containing 1% SDS and 2-mercaptoethanol, followed by serial dilution in five 2-fold steps with dilution lysis buffer. The diluted lysates were arrayed onto nitrocellulose-coated slides (Grace Bio-Labs) using the Quanterix (Aushon) 2470 Arrayer (Quanterix Corporation), resulting in a total of 5808 spots per slide. These spots included serial dilutions of standard lysates, as well as positive and negative controls prepared from mixed cell lysates and dilution buffer, respectively.

Each slide underwent probing with a validated primary antibody and a biotin-conjugated secondary antibody. The validation process for antibodies used in reverse-phase protein array (RPPA) is detailed on the RPPA Core website (https://www.mdanderson.org/research/research-resources/core-facilities/functional-proteomics-rppa-core/antibody-information-and-protocols.html). Signal amplification was achieved using the Agilent GenPoint staining platform (Agilent Technologies), and the signal was visualized through a DAB colorimetric reaction. Slides were scanned using a Huron TissueScope (Huron Digital Pathology) and quantified with Array-Pro Analyzer software (Media Cybernetics) to obtain spot intensity data.

### RPPA data analysis

Relative protein levels for each sample were determined using RPPA SPACE, a software tool developed by the MD Anderson Department of Bioinformatics and Computational Biology (https://bioinformatics.mdanderson.org/public-software/rppaspace).^69^ This tool fits a logistic model to each dilution curve, utilizing all samples on a slide. The fitting process plots observed and fitted signal intensities against the log2 concentration of proteins for diagnostic purposes. Protein concentrations were then normalized for protein loading by calculating a correction factor through median-centering across samples for all antibody experiments and across antibodies for each sample. Normalization across RPPA sets was performed using replicates-based normalization as described.^70–71^

### RT-qPCR

Total RNA was extracted from the sorted cells as specified previously. The concentration and purity of RNA were determined using a NanoDrop spectrophotometer. Reverse transcription was performed using the iScript™ cDNA Synthesis Kit (Bio-Rad Laboratories, Inc.), and the reaction was carried out in a thermal cycler according to the manufacturer’s instructions. The resulting cDNA was diluted and used as a template for qPCR, which was performed in a 384-well plate format with triplicate wells for each condition. The qPCR was performed using either the TaqMan Fast kit (Thermo) or the PowerTrack SYBR Green kit (Thermo). The following TaqMan primer/probe sets were used: Amy2a3 (Mm02342486_mH), Cpa1 (Mm00465942_m1), Mist1 (Mm00627532_s1), Ptfla (Mm00479622_m1), Hnf6a (Hs00413554_m1), Krt19 (Mm00492980_m1), Pdx1 (Mm00435565_m1), Sox9 (Mm00448840_m1).

### Lysate preparation

Pancreata were excised, washed quickly in 5mL of ice-cold HBSS to remove excess blood and contaminating lymph nodes. Then tissue was put into a 1.5 mL microtube, and then submerged into liquid nitrogen for snap freezing. Frozen tissue was crushed into a powder, and samples were resuspended in 500 μl of RIPA+PIC solution (PIC diluted 1 to 100) and incubated on ice for 15-20 min with intermediate vortex every 5 min. Samples were transferred into cold 1.5 mL tube and centrifuged at 10,000g for 10min at 4°C. Supernatant was transferred to a new tube. Protein concentration quantifications were performed using Pierce BCA Protein Assay Reagent (ThermoFisher Scientific, Cat# 23235) following the supplier protocol.

### Western blotting analysis and active Ras/Rac1 pull-down assays

Standard techniques were employed for immunoblotting. Protein lysates were separated on 4%–12% Bis-Tris NuPage gels (Life Technologies). Western blots were probed with the following antibodies: Anti-Ras Antibody (Active Ras Pull-Down and Detection Kit, ThermoFisher Scientific, 16117), Akt (pan) Rabbit mAb (C67E7, Cell Signaling Technology, 4691S), Phospho Akt (Ser473) Rabbit mAb (193H12, Cell Signaling Technology, 4058S), Anti-beta Actin antibody (Abcam, ab8227), P44/42 MAPK (Erk1/2) Mouse mAb (L34F12, Cell Signaling Technology, 4696S), Phospho-p44/42 MAPK (Erk1/2) Rabbit Monoclonal Antibody (Thr202/Tyr204, D13.14.4E, Cell Signaling Technology, 4370S), Anti-Vinculin antibody (Abcam, ab129002), Anti-Rac1 Antibody (Active Rac1 Pull-Down and Detection Kit, ThermoFisher Scientific, 16118), Vav1 Antibody (Cell Signaling Technology, 2502S), GAPDH Rabbit mAb (D16H11, Cell Signaling Technology, 5174S), and Rabbit anti-TIAM1 Antibody (Bethyl Laboratories, A300-099A). The levels of Ras-GTP were determined using the Active Ras Pull-Down and Detection Kit (ThermoFisher Scientific, 16117) following the manufacturer’s instructions. Similarly, the Active Rac1 Pull-Down and Detection Kit (ThermoFisher Scientific, 16118) was used for determining Rac1-GTP levels according to the manufacturer’s instructions. Detection was performed using chemiluminescence, and images were captured using a Bio-Rad ChemiDoc Imaging System.

### Quantification and statistical analysis

The number of animals is based on feasibility considerations and our interest in observing a relatively large effect size. In addition, a power analysis was performed with a desired power of 0.8, α= 0.05, and a literature search was performed to find expected averages and standard deviations based on similar protocols. Tests for differences between two groups were performed using two-tailed unpaired Student t test or two-tailed Mann–Whitney test as specified in the figure legends. P values were considered significant if less than 0.05. All graphs depict mean ± SEM unless otherwise indicated. Asterisks used to indicate significance correspond with *, P < 0.05; **, P < 0.01; ***, P < 0.001; ****, P < 0.0001. GraphPad Prism 9 or Prism 10 (GraphPad Software) was used for statistical analysis of experiments, data processing, and presentation.

## DATA AVAILABILITY STATEMENT

All high-throughput sequencing data, both raw and processed files, have been deposited in NCBI’s Gene Expression Omnibus and are publicly available. Accession numbers are listed in the key resources table. Microscopy data reported in this paper will be shared by the corresponding authoer upon request. This paper does not report original code. Mouse lines (Ptf1a-tdTomato) generated in this study have been deposited to The Jackson Laboratory. Further information and requests for resources and reagents should be directed to and will be fulfilled by the corresponding author, Rohit Chandwani.

