## Supplementary Figures for "Molecular dynamics driving phenotypic divergence among KRAS mutants in pancreatic tumorigenesis"

**SUPPLEMENTARY INFORMATION**

**Figure S1. Injury induces lineage reversion to a dedifferentiated cell state. (A)** Schematic representation of how the tdTomato-2A-NSF cassette was inserted into the Ptf1a locus using recombinase-mediated cassette exchange to generate a novel lineage tracing mouse model (i.e., Ptf1a-tdTomato). **(B)** Whole-organ fluorescent imaging of adult Ptf1a-tdTomato mouse pancreas. Representative images display normal pancreatic morphology with tdTomato expression in the pancreas of both heterozygous (Ptf1a-tdT/+) and homozygous (Ptf1a-tdT/tdT) mice. **(C)** Immunofluorescence staining for TdTomato, GFP and DAPI markers on collagenase digested pancreas isolated from an adult $Ptf1a^{tdT/+}$; $Pdx1^{GFP/+}$ mouse pancreas. **(D)** Immunofluorescence staining for DAPI, tdTomato, E-Cadherin, and C-peptide on $Ptf1a^{tdT/+}$ mouse pancreas. Scale bar, 100 µm. **(E)** Flow cytometric analysis of embryonic day 10.5 (e10.5) and embryonic day 15.5 (e15.5) pancreatic progenitors from Ptf1a-tdTomato mice, as well as postnatal day 60 (p60) C57BL/6 mice stained with Cd49f and Cd133. **(F)** Barplot illustrating fold-induction of representative marker genes measured using RT-qPCR on FACS-sorted adult exocrine cells (acinar and ductal) derived from p60 C57BL/6 mice in E. **(G)** Barplot of RNA-seq data visualizing the number of transcripts up- and down-regulated across sorted pancreatic cell populations Thresholds: absolute log2(fold change) > 0; padj < 0.01. **(H)** Schematic representation of lineage-traced mouse model, and representative hematoxylin and eosin staining of caerulein-treated and saline-treated pancreata. Representative images of n=4-5 mice per condition. Scale bar, 100 µm. **(I)** Principal Component Analysis (PCA) of RNA-seq data from injured (ADM) and normal (acinar, ductal, early progenitors, late progenitors) pancreas samples. **(J)** Euclidean distance analysis of ADM cells relative to acinar, ductal, as well as early and late progenitor cells. **(K)** Gene Set Enrichment Analysis (GSEA) of ADM versus control acinar cells. GSEA results for significantly enriched gene sets are displayed.

**Figure S2. Development and implementation of gene signatures to contextualize metaplasia. (A-E)** Heatmap of RNA-seq data illustrating gene expression differences between **(A)** exocrine (acinar, ductal) and progenitor (early, late) cell populations; **(B)** acinar UP and Acinar DOWN; **(C)** ductal UP and ductal down; **(D)** early progenitor UP and early progenitor DOWN; and **(E)** late progenitor UP and late progenitor DOWN. Representative genes are highlighted alongside the heatmap. Thresholds: absolute log2(fold change) > 0; padj < 0.01. Each column is a biological replicate mouse. **(F)** Barplot of RNA-seq data visualizing the number of transcripts up- and down-regulated across all pairwise contrasts between acinar, ductal, early progenitor, late progenitor, ADM cells. Thresholds: absolute log2(fold change) > 1; padj < 0.01. **(G)** Heatmap of RNA-seq data visualizing the expression of intersecting genes from two comparisons: ADM versus acinar & ductal versus late progenitor. The expression levels are from the samples listed (ADM, late progenitor, and ductal cells). Thresholds: absolute log2(fold change) > 1; padj < 0.01. Each column is a biological replicate mouse. **(H)** Dot plot illustrating z-score of gene abundance for intersecting genes from two comparisons: ADM versus acinar & ductal versus late progenitor. Four distinct groups of expression dynamics are presented (Group 1: 121 genes, Group 2: 643 genes, Group 3: 791 genes, Group 4: 66 genes).

**Figure S3. Molecular alterations induced by *$Kras^{G12D}$*. (A)** Gene Ontology (GO) term enrichment analysis of RNA-seq data comparing tdTomato+ cells isolated from WT to tdTomato+ cells collected from $MK^{G12D}T$ mice 7 weeks of mutant Kras activation. **(B)** Tornado plots visualizing ATAC-seq data of previously described KRAS-induced DARs from GSE132330 (Alonso-Curbelo et al., 2021). Left plot: normal acinar and $Kras^{G12D}$ samples from Alonso-Curbelo et al., 2021. Right plot: normal acinar and 7 week $Kras^{G12D}$ samples from the current study. Thresholds: log2FC ≥ 0.58; FDR ≤ 0.1 (as defined by Alonso-Curbelo et al, 2021).

**Figure S4. *$Kras^{G12R}$* drives robust transcriptional rewiring early after activation. (A)** PCA of RNA-seq data for the top 2000 most variable genes in tdTomato+ cells isolated from WT and tdTomato+ cells isolated from $MK^{G12D}T$, $MK^{G12Vgeo}T$, and $MK^{G12R}T$ mice after 10 days of mutant Kras activation. **(B)** Bhattacharyya distance analysis of the top 2000 most variable genes in tdTomato+ cells isolated from WT (MT), $MK^{G12D}T$, $MK^{G12Vgeo}T$, and $MK^{G12R}T$ mice after 10 days of mutant Kras activation. **(C)** Dot plot illustrating z-score of gene abundance for DEGs identified from normal acinar cells versus $MK^{G12D}T$ at 7 weeks (N = 551). Distinct groups of expression dynamics are presented; only the groups containing at least 25 genes are included. The red and blue lines indicate expression patterns at 10 days and 6 weeks, respectively. **(D)** GSEA performed on RNA-seq data using adult and developing pancreas benchmark gene sets, independently comparing tdTomato+ cells isolated from WT to tdTomato+ cells collected from $MK^{G12D}T$, $MK^{G12Vgeo}T$, and $MK^{G12R}T$ mice after 10 days.

**Figure S5. An emergent hierarchy between mutant Kras alleles in the context of injury. (A)** Immunofluorescence for CPA1 / CK19 and Sox9 / tdTomato staining of pancreas sections collected from acutely injured MT, $MK^{G12D}T$,

MK$^{G12Vgeo}$T, and MK$^{G12R}$T mice 2 days post-injury. Representative images of n=2-4 mice per condition. Scale bar, 100 µm. **(B)** Dot plot illustrating z-score of gene abundance for DEGs identified by comparing normal acinar cells versus injured MK$^{G12D}$T cells collected 2 days post-injury (N = 5424; Thresholds: absolute log2(fold change) > 1; padj < 0.01). Distinct groups of expression dynamics are presented across genotypes; only the groups containing at least 25 genes are included. **(C)** Heatmap of DEGs (encoding transcription factors, cytokines, and kinases) identified by comparing tdTomato+ cells collected from acutely injured MK$^{G12D}$T and MK$^{G12R}$T mice 2 days post-injury. Thresholds: padj < 0.01. **(D)** GSEA performed on RNA-seq data comparing tdTomato+ cells collected from MK$^{G12D}$T mice to both MK$^{G12R}$T (top) and MK$^{G12Vgeo}$T (bottom) mice 2 days post-acute injury. **(E)** GO analysis performed on RNA-seq data comparing tdTomato+ cells collected from MK$^{G12D}$T mice to both MK$^{G12R}$T (top) and MK$^{G12Vgeo}$T (bottom) mice 2 days post-acute injury. **(F)** HOMER analysis depicting top motif enrichments when comparing tdTomato+ cells collected from both MK$^{G12D}$T mice and MK$^{G12R}$T mice 2 days post-acute injury. **(G)** Hematoxylin and eosin staining of MT (WT), MK$^{G12D}$T, MK$^{G12Vgeo}$T, and MK$^{G12R}$T mouse pancreas sections collected 21 days post-acute injury. Representative images of n=2-9 mice per condition. Scale bar, 100 µm. **(H)** Histologic quantifications of normal acinar tissue, ADM, and PanIN lesions in pancreata collected from MT, MK$^{G12D}$T, MK$^{G12Vgeo}$T, and MK$^{G12R}$T mice 21 days post-acute injury.

**Figure S6. Molecular differences between Kras mutants in the context of prolonged injury. (A)** GSEA performed on RNA-seq data comparing tdTomato+ cells isolated from WT without injury to tdTomato+ cells collected from both MK$^{G12R}$T mice (left) or MK$^{G12V}$T mice (right) 3 weeks post-acute injury. **(B)** GSEA performed on RNA-seq data comparing tdTomato+ cells collected from MK$^{G12D}$T mice to both MK$^{G12R}$T mice (left) or MK$^{G12V}$T mice (right) 3 weeks post-acute injury. **(C)** GO analysis performed on RNA-seq data comparing tdTomato+ cells collected from MK$^{G12D}$T mice to both MK$^{G12R}$T mice (top) or MK$^{G12V}$T mice (bottom) 3 weeks post-acute injury. Top 5 terms are shown. **(D)** Representative tornado plots visualizing ATAC-seq data from the current study: normal acinar cells, MT, MK$^{G12D}$T, MK$^{G12Vgeo}$T, and MK$^{G12R}$T mice 3 weeks after prolonged injury. DARs identified by comparing PDAC vs. WT samples from GSE132330 (Alonso-Curbelo et al., 2021). Thresholds: log2FC ≥ 0.58; FDR ≤ 0.1. Each column is a biological replicate mouse.

**Figure S7. Kras$^{G12R}$ and Kras$^{G12V}$ feature impaired EGFR signaling. (A)** Cytokine Signaling Analyzer (CytoSig) output reflecting enriched cytokine signaling pathways for the indicated conditions. Each column reflects the transcriptional contribution of an individual mouse. **(B)** Dot plot indicating differential expressed cytokines between G12D and G12R (padj < 0.01) in either acute (blue circles) or prolonged (orange circles) injury. The top 10 cytokines enriched in G12D with acute injury are highlighted in black, and Areg/Ereg are highlighted in red. **(C-D)** Cytokine array indicating relative enrichment of the indicated cytokines in the G12D, G12V, and G12R pancreas compared with WT, where all tissues are harvested 2 days after acute injury. **(E)** Brightfield and fluorescent microscopy images of acinar cells treated with either vehicle (top half) or recombinant human TGFα (bottom half) to induce *in vitro* ADM. Scale bar, 500 µm. **(F)** Quantification of mean object area of *in vitro* ADM as in **(E)**; each point represents a well of cells belonging to the indicated treatment and genotype. **(G)** Western blot showing EGFR Y1068 phosphorylation, total EGFR, and vinculin from bulk pancreas tissue lysate extracted from MT, MK$^{G12D}$T, MK$^{G12Vgeo}$T, and MK$^{G12R}$T 2 days following acute injury. **(H)** Quantification of EGFR signal normalized to vinculin, as per data shown in **(G)**. **(I)** Quantification of phospho-EGFR signal normalized to total EGFR, as per data shown in **(G).**

**Figure S8. Signaling defects in the Kras$^{G12R}$ mutant pancreas. (A)** Western blot analysis showing total Ras and Ras-GTP levels in MT, MK$^{G12D}$T, MK$^{G12Vgeo}$T, and MK$^{G12R}$T mice 7 weeks post-tamoxifen injection. Quantification of Ras-GTP levels relative to total Ras. **(B)** Quantification of p-Erk signal normalized to Erk, as per data shown in Figure 6C. **(C)** Western blot analysis of total Erk and phosphorylated Erk (p-Erk) levels in MT, MK$^{G12D}$T, MK$^{G12Vgeo}$T, and MK$^{G12R}$T bulk pancreatic lysate collected 2 days after acute injury. Quantification of p-Erk levels relative to total Erk. **(D)** Western blot analysis of total Erk, p-Erk and vinculin levels in MT, MK$^{G12D}$T, MK$^{G12Vgeo}$T, and MK$^{G12R}$T bulk pancreatic lysate collected 3 weeks post-prolonged injury. Quantification of p-Erk levels relative to total Erk. **(E-F)** Quantification of Vav1 signal normalized to GAPDH, as per data shown in **(E)** Figure 6H and **(F)** Figure 6J**. (G)** Western blot showing Vav1 and GAPDH expression levels from sorted tdTomato+ cells collected from MT, MK$^{G12D}$T, MK$^{G12Vgeo}$T, and MK$^{G12R}$T mice [3 weeks after prolonged injury]. **(H)** Western blot analysis of Vav1 and GAPDH levels in MT, MK$^{G12D}$T, MK$^{G12Vgeo}$T, and MK$^{G12R}$T bulk pancreatic lysate 2 days after acute injury. GAPDH was used as a loading control. Quantification of Vav1 relative to GAPDH.

### Figure S1. Injury induces lineage reversion to a dedifferentiated cell state

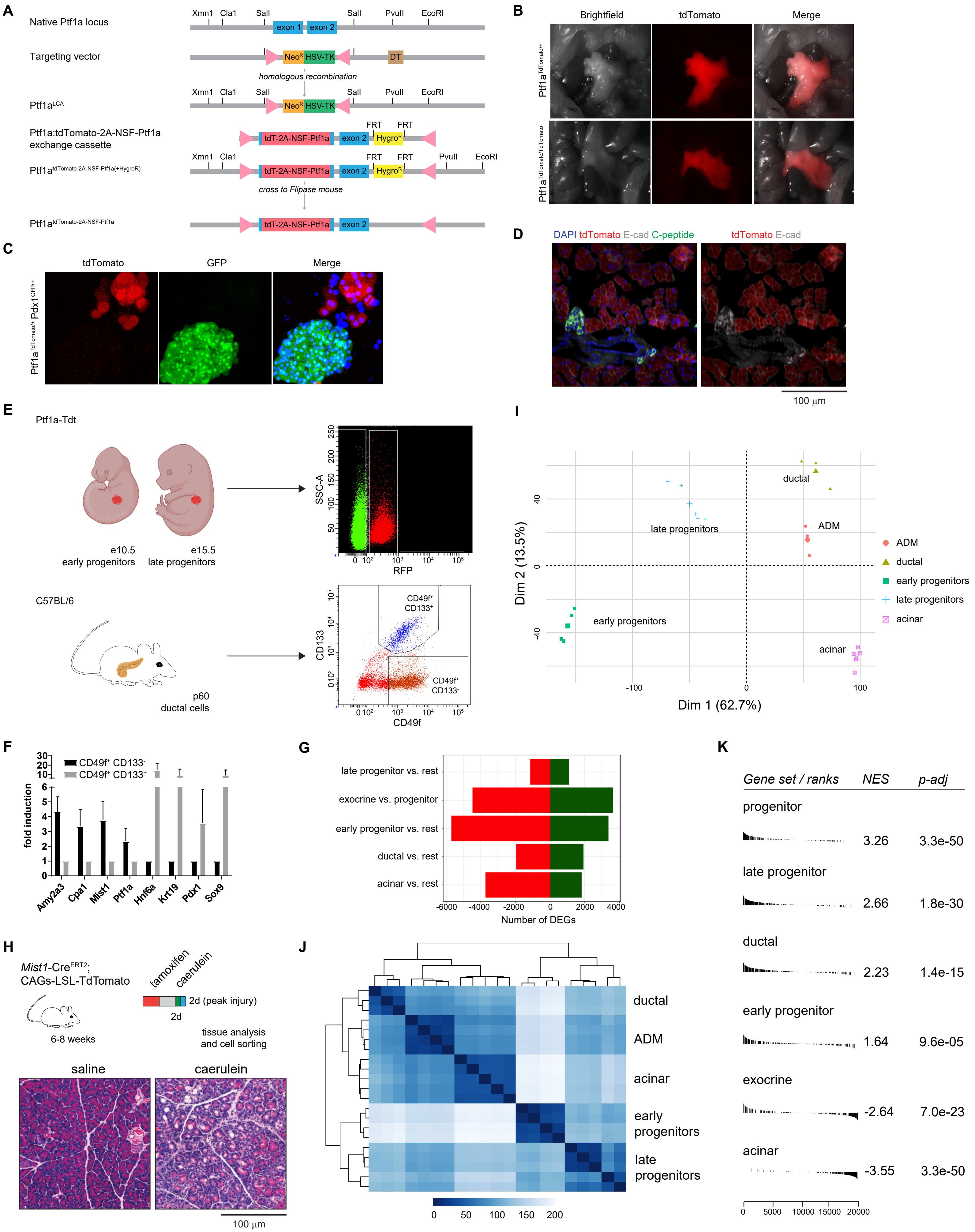

### Figure S2. Development and implementation of gene signatures to contextualize metaplasia.

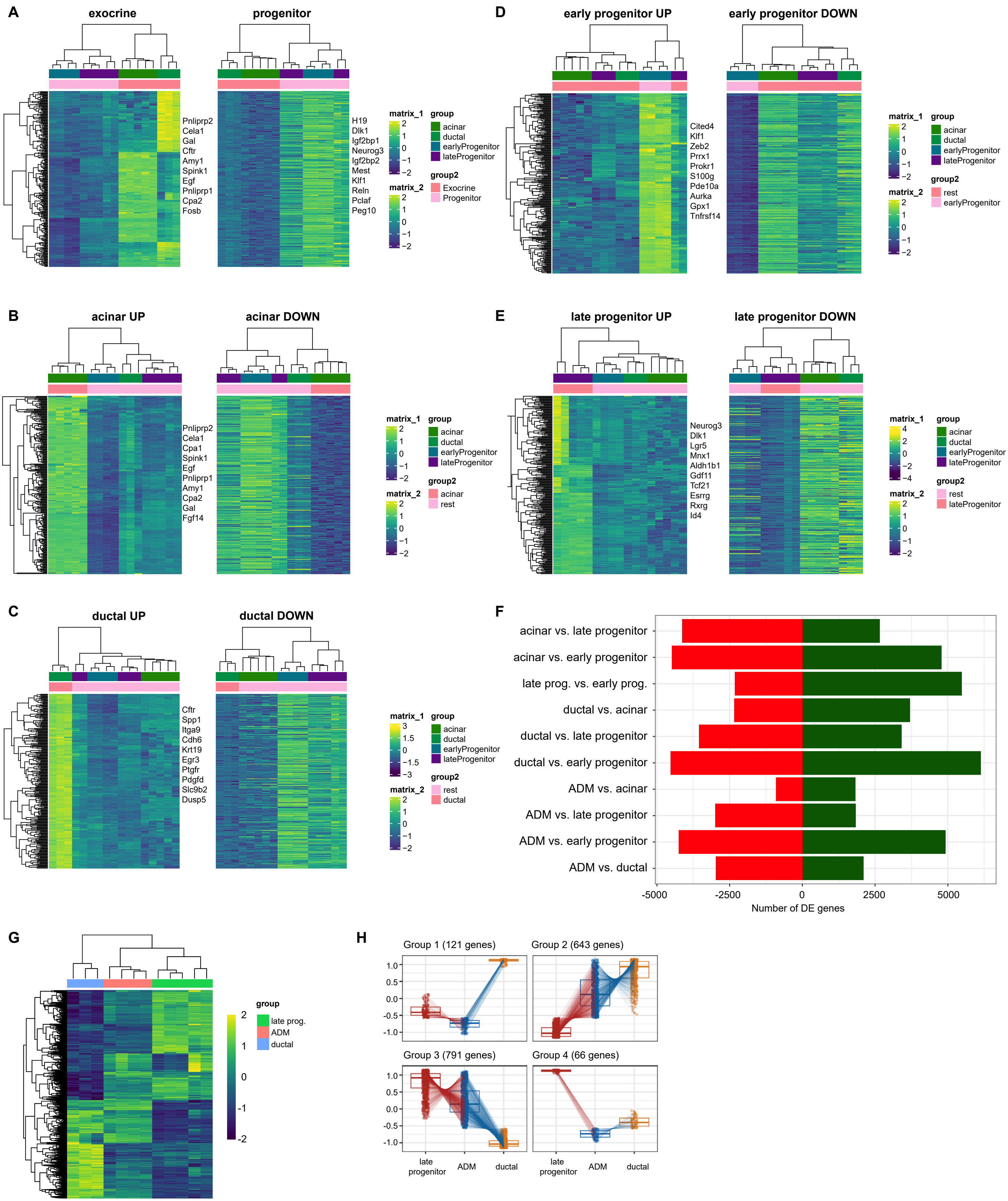

**Figure S3. Molecular alterations induced by *Kras*$^{G12D}$.**

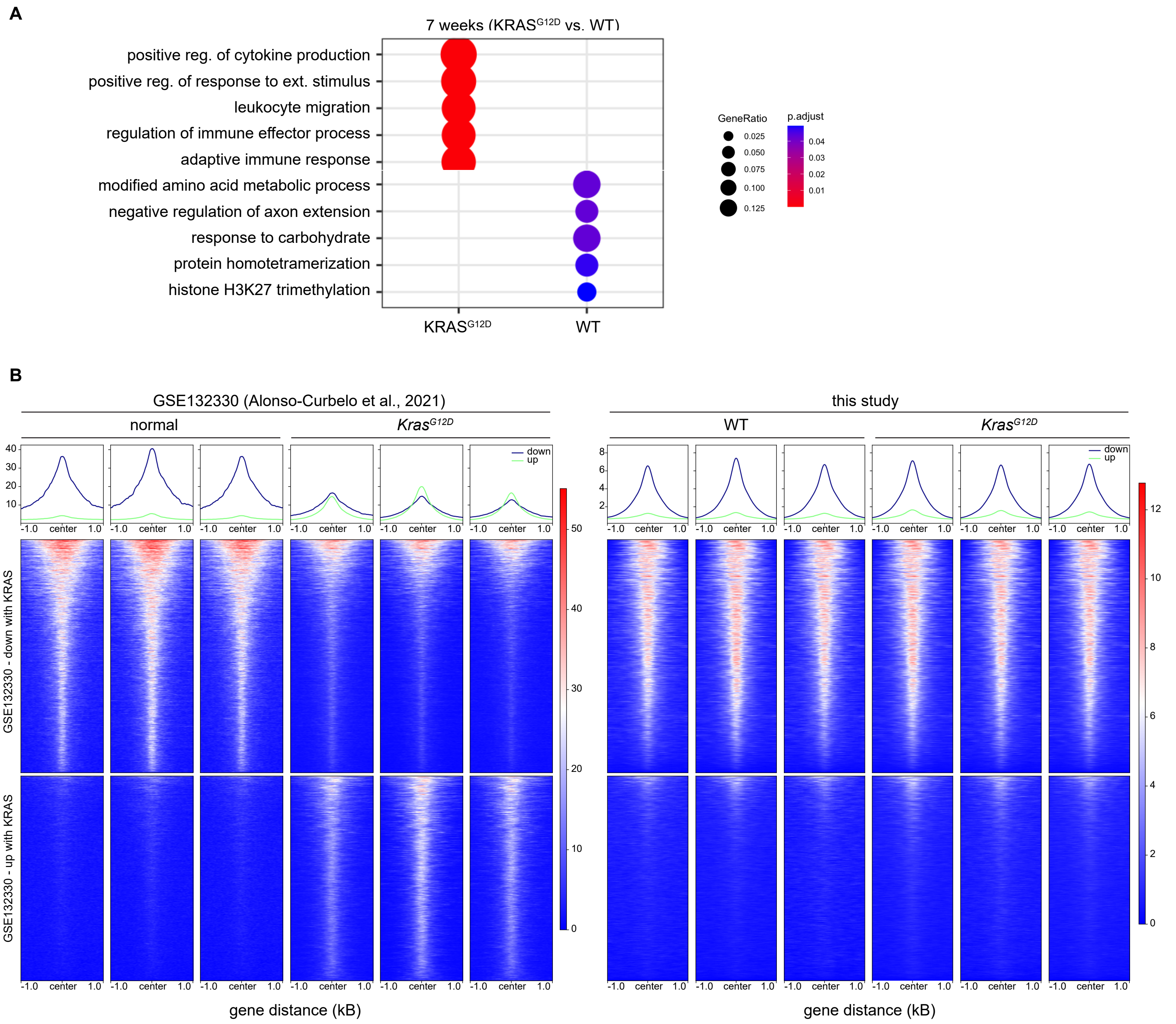

**Figure S4. *Kras*$^{G12R}$ drives robust transcriptional rewiring early after activation.**

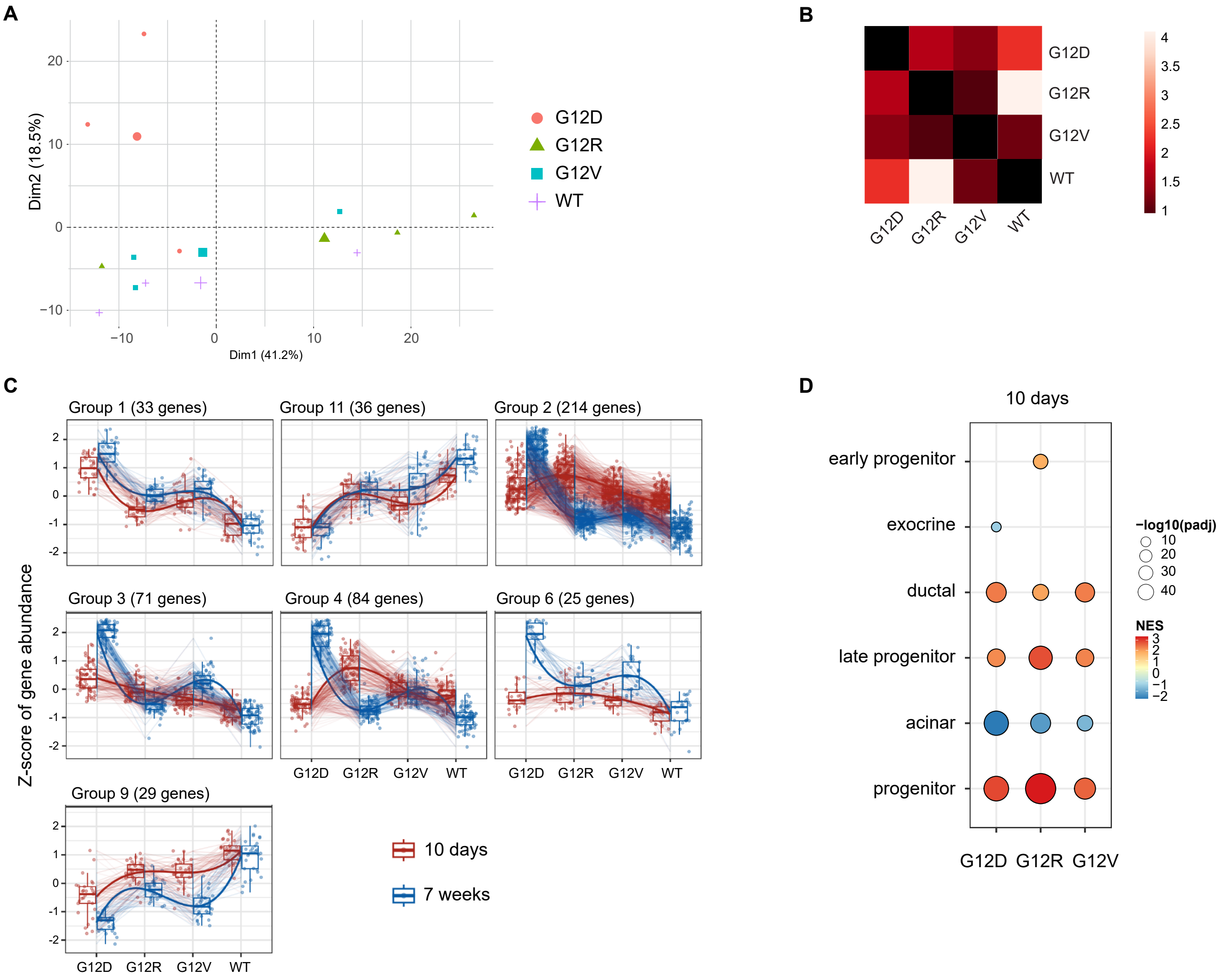

**Figure S5. An emergent hierarchy between mutant *Kras* alleles in the context of injury.**

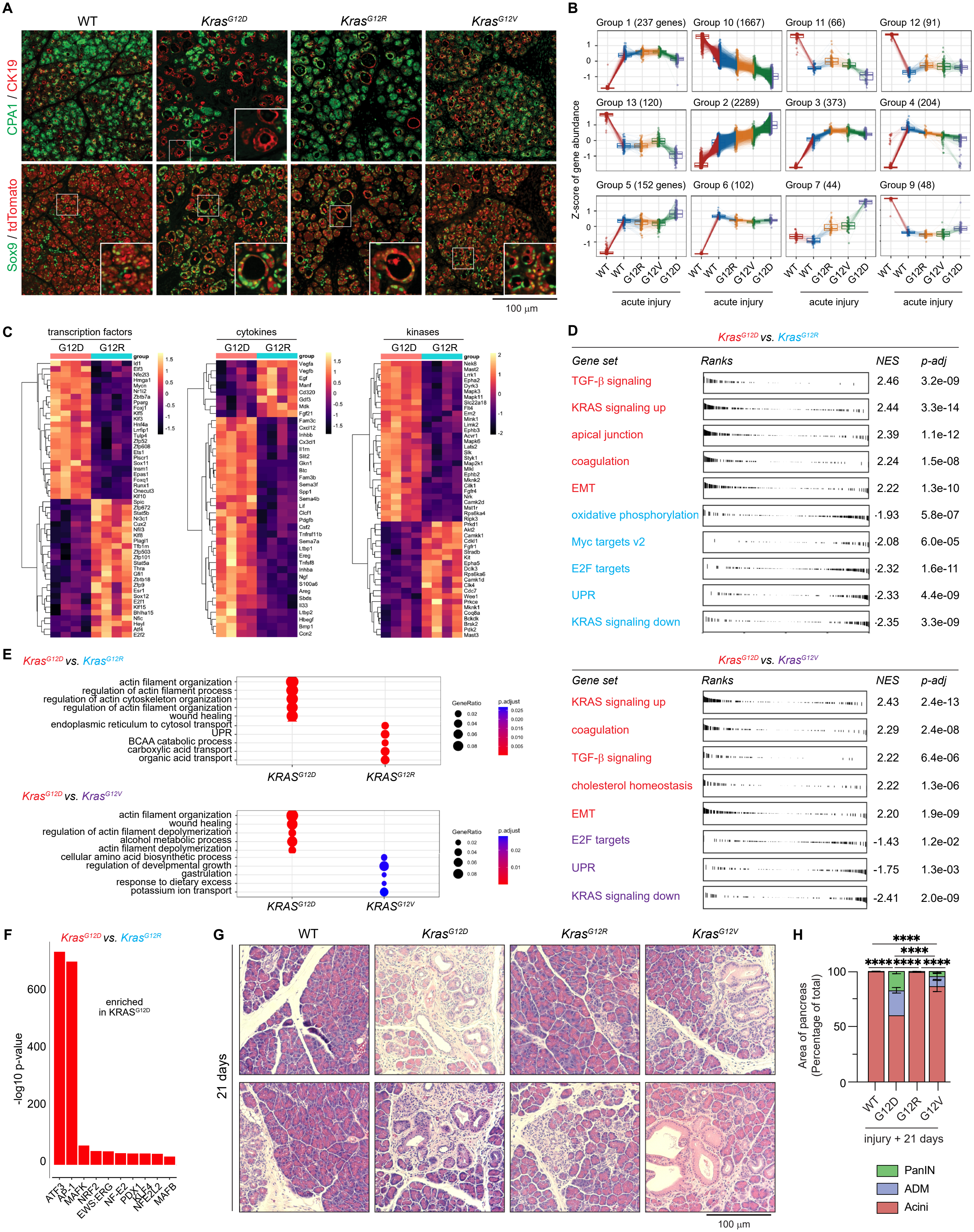

#### Figure S6. Molecular differences between *Kras* mutants in the context of prolonged injury.

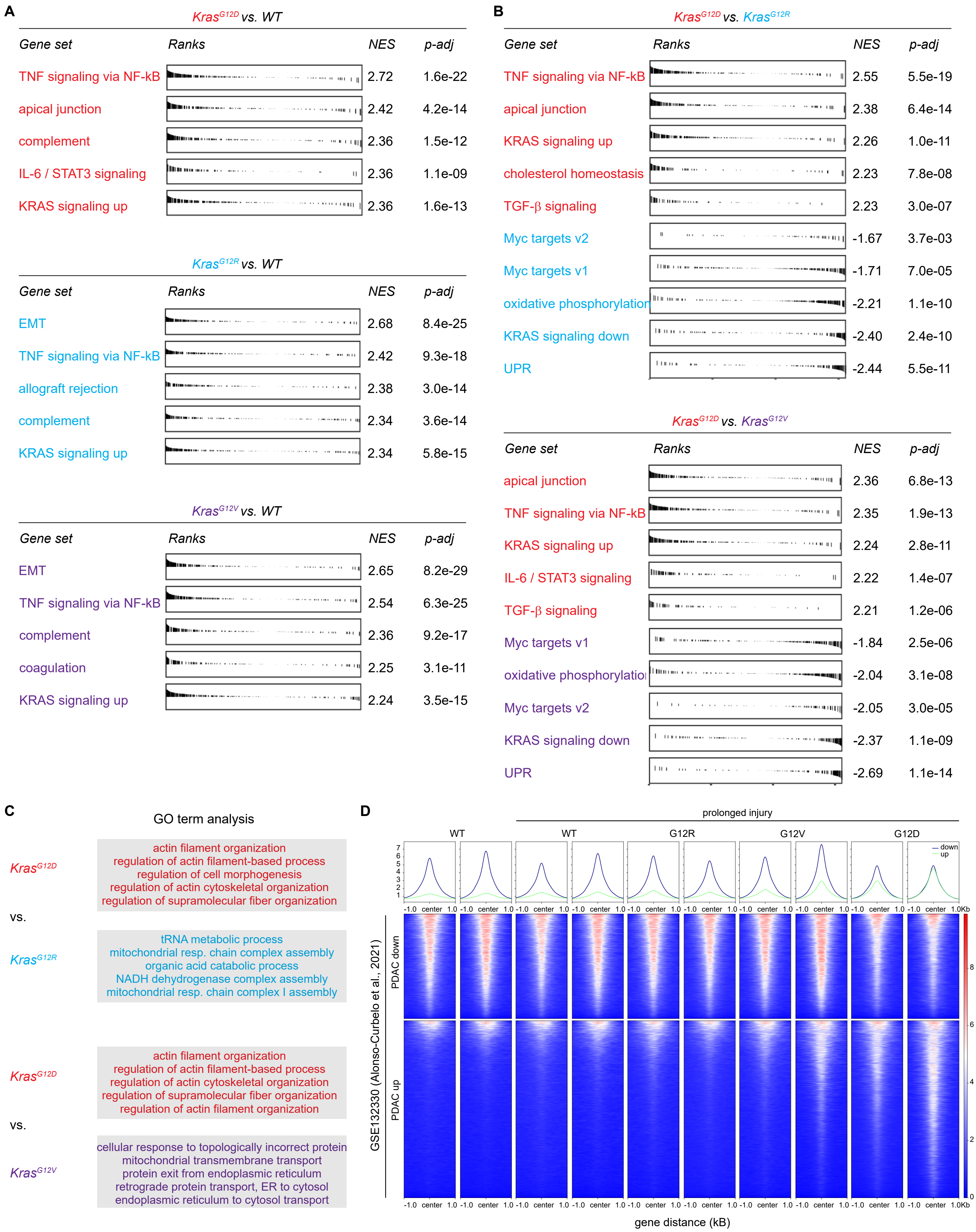

#### Figure S7. *Kras*$^{G12R}$ and *Kras*$^{G12V}$ feature impaired EGFR signaling

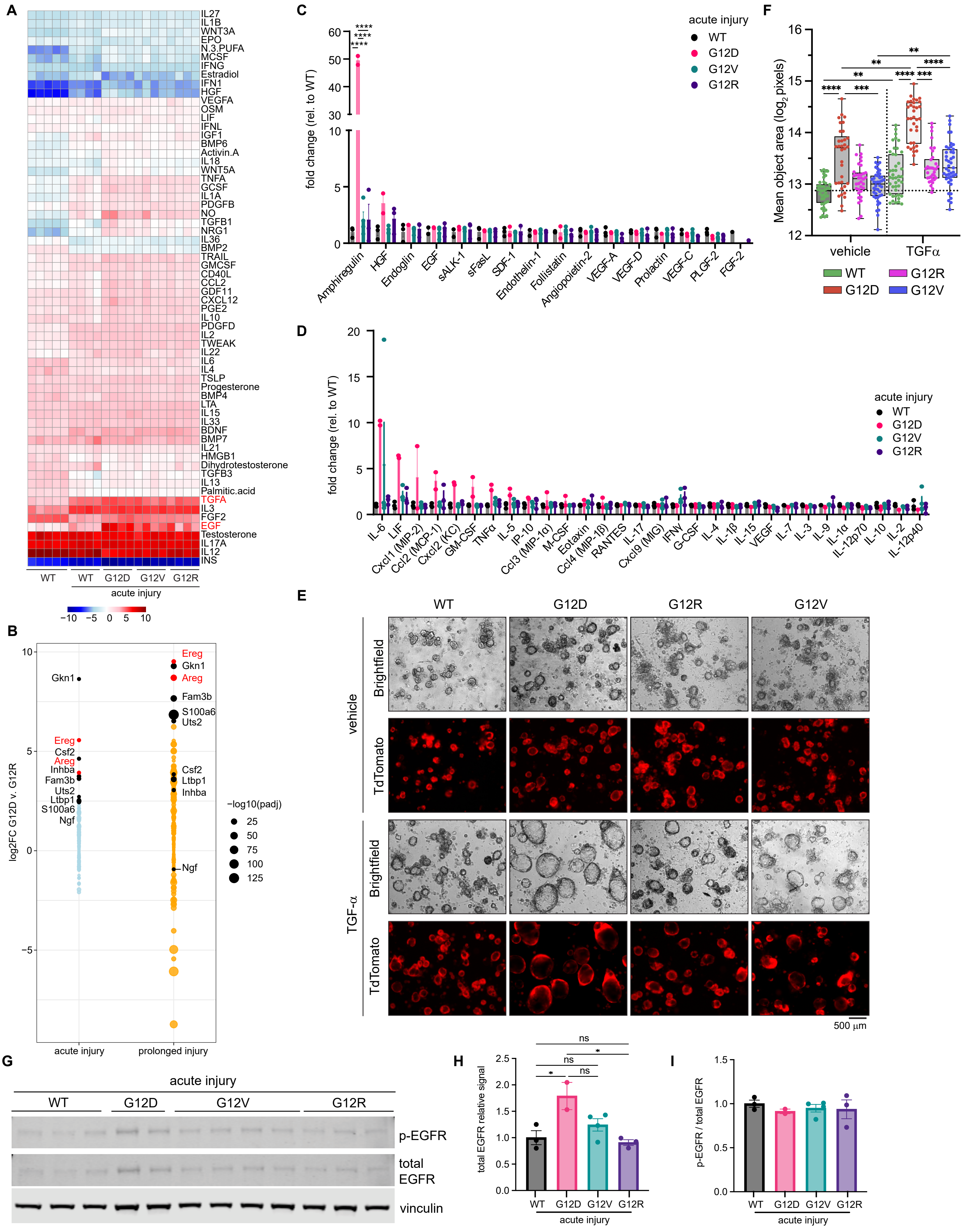

**Figure S8. Signaling defects in the *Kras*$^{G12R}$ mutant pancreas.**

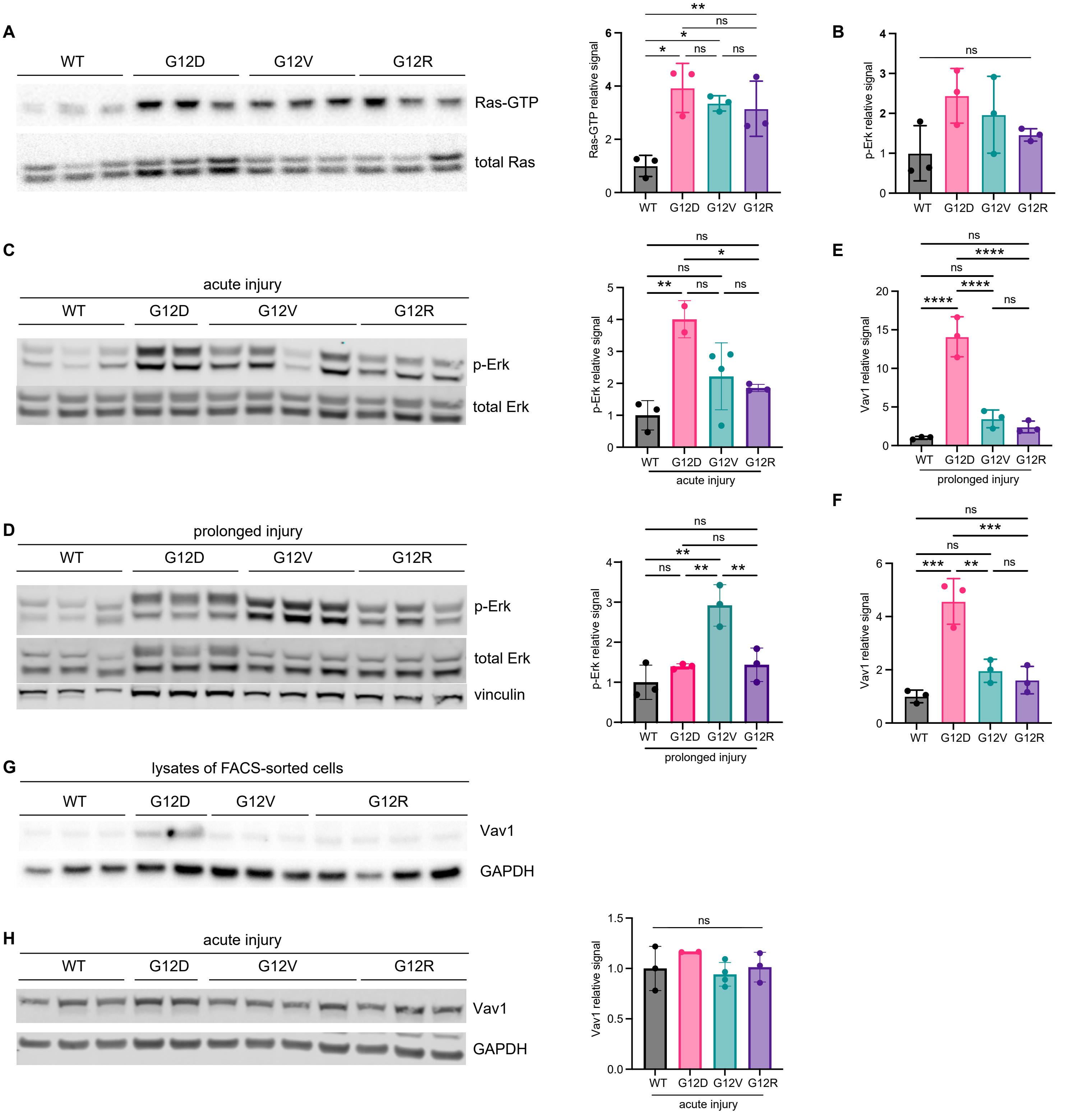
